# Four ways to fit an ion channel model

**DOI:** 10.1101/609875

**Authors:** M. Clerx, K.A. Beattie, D.J. Gavaghan, G.R. Mirams

## Abstract

Computational models of the cardiac action potential are increasingly being used to investigate the effects of genetic mutations, predict pro-arrhythmic risk in drug development, and to guide clinical interventions. These safety-critical applications, and indeed our understanding of the cardiac action potential, depend on accurate characterisation of the underlying ionic currents. Four different methods can be found in the literature to fit ionic current models to single-cell measurements: (Method 1) fitting model equations directly to time constant, steady-state, and I-V summary curves; (Method 2) fitting by comparing simulated versions of these summary curves to their experimental counterparts; (Method 3) fitting to the current traces themselves from a range of protocols; and (Method 4) fitting to a single current trace from an information-rich voltage clamp protocol. We compare these methods using a set of experiments in which hERG1a current from single Chinese Hamster Ovary (CHO) cells was characterised using multiple fitting protocols and an independent validation protocol. We show that Methods 3 and 4 provide the best predictions on the independent validation set, and that the short information-rich protocols of Method 4 can replace much longer conventional protocols without loss of predictive ability. While data for Method 2 is most readily available from the literature, we find it performs poorly compared to Methods 3 and 4 both in accuracy of predictions and computational efficiency. Our results demonstrate how novel experimental and computational approaches can improve the quality of model predictions in safety-critical applications.

**Statement of Significance:** Mathematical models have been constructed to capture and share our understanding of the kinetics of ion channel currents for almost 70 years, and hundreds of models have been developed, using a variety of techniques. We compare how well four of the main methods fit data, how reliable and efficient the process of fitting is, and how predictive the resulting models are for physiological situations. The most widely-used traditional approaches based on current-voltage and time constant-voltage curves do not produce the most predictive models. Short, optimised experimental voltage clamp protocols can be used to create models that are as predictive as ones derived from traditional protocols, opening up possibilities for measuring ion channel kinetics faster, more accurately and in single cells. As these models often form part of larger multi-scale action potential and tissue electrophysiology models, improved ion channel kinetics models could influence the findings of thousands of simulation studies.

## INTRODUCTION

Computational models of ionic currents have been used to understand the formation of the action potential (AP) (1–3), the effects of genetic mutations (4), and the action of drugs on the rhythm of the heart (5). By fitting a mathematical model to experimentally measured currents we can learn about the kinetics of the underlying ion channels, and how they might be altered in pathological conditions or the presence of a drug. Being able to do this rapidly and accurately is becoming increasingly important as cardiac modelling progresses from a tool for laboratory investigation to a tool for prediction in safety-critical applications in drug development and clinical risk assessment (6–8).

Several methods have been proposed to fit (parameterise) models of ionic currents, covering a range including: pen-and-paper methods of the 1950s; detailed mathematical analysis specific to Hodgkin-Huxley models; and numerical ‘black-box’ optimisation. Data from several sources has been used, including whole-cell (aggregate) currents, single-channel currents (9), action potential recordings (10), dynamic-clamp experiments (11), gating currents (12), and measurements of fluorophores whose visibility varies with channel state (13).

In this manuscript, we focus on the common problem of fitting a set of predetermined model equations to voltage-dependent whole-cell ionic currents recorded under voltage clamp. We describe four methods of fitting, each representative of a wider class of methods used in the electrophysiology literature. Data from a previous study is used, where currents were recorded at room temperature in nine Chinese Hamster Ovary (CHO) cells stably expressing hERG1a (14), and where each cell was subjected to a series of protocols used for fitting and an independent protocol used for validation. We apply all four methods to each cell, creating four model parameterisations, and then evaluate their ability to predict the current measured in the validation protocol. Finally, we discuss some of the reasons behind the observed difference in performance, and compare the methods’ repeatability and time-efficiency.

The first publication describing and fitting of a model of ionic currents was by Hodgkin and Huxley (1). Using the idea that ionic conductance depends on some part of the membrane being in one of two states, they postulated a simple two-state system with a forward and a backward transition rate from which a time constant and a steady state could be deduced. They then set about designing protocols to measure (or approximate) these time constants and steady states at several voltages, fitted curves through these measurements, and used the resulting (phenomenological) equations to encode their transition rates. Similar methods of fitting the equations directly to experimentally derived points — using mathematical analysis, least-squares fitting, or manual adjustment — were employed by subsequent modellers (15–24).

In this work, we describe a version of this method, and refer to it and similar methods as ‘*Method 1*’. It is important to note that these methods rely on particular assumptions: (1) that the underlying biophysics is accurately described by one or more independent gating processes, each with their own steady state and time constant variables; and (2) that an experimental method and analysis procedure will yield accurate values for these variables at a range of physiologically relevant voltages. The first assumption applies only to Hodgkin-Huxley style models — there are no such analytic expressions available to fit when using Markov models, making Method 1 unusable for Markov models. Testing if the second assumption is violated can be done using simulated experiments, but proving it holds is challenging. A detailed mathematical analysis of two protocols was performed by Beaumont et al. (25), who showed that steady states can be obtained from peak currents only if there is a large difference in the time constants of the system, and that once these are known the time constants can be estimated using time-to-peak measurements or by fitting exponential curves. In a follow-up paper they presented an iterative procedure to estimate the steady-states for systems with more similar (‘non-separable’) time constants (26). In Wang and Beaumont (27) this analysis is taken further and a method is derived to estimate steady-state equations and time constants simultaneously and further improve the results (the idea to fit both parameters together had also been used previously by Ebihara and Johnson (28)). Willms et al. (29) provided another mathematical analysis of fitting two-state Hodgkin-Huxley models, and conclude that separate estimation of time constants and steady states (‘the disjoint method’) can lead to poor results. A further critique of Method 1 is given by Lee et al. (30), who use mathematical analysis and simulation to point out errors arising for non-separable time constants, and introduce methods aimed to combat this effect.

Despite these critiques, Method 1 remains highly popular due to its simplicity. Estimates for time constants and steady states are easily obtainable from the literature, which is a distinct advantage when multiple currents must be considered e.g. when creating full action potential models (31–34).

An alternative to Method 1, but using the same data points, is to simulate the voltage protocols and the subsequent analysis to obtain a set of simulated steady states and time constants to match the experimental ones. The parameters used in the simulation can then be updated in an iterative fashion (manually or using numerical optimisation) until the simulated values match the experimental ones. This procedure allows both Hodgkin-Huxley and Markov models to be fit to published time-constant and steady-state data (although a unique/identifiable fit is not guaranteed). If the modeller uses the same analysis method as was used on the experimental data, any imperfections in the approximations of the steady states and time constants are replicated in the simulation, so that good results may be obtainable even when Method 1 approximations are poor (e.g. if time constants are non-separable). In this manuscript, we use the name ‘*Method 2*’ to describe such methods, and define and analyse a version of this method based on numerical optimisation.

Like Method 1, Method 2 is prevalent in the AP-modelling literature (35–42). Further users include Moreno et al. (43), who mention the need to use published data as the reason to prefer Method 2, as well as Perissinotti et al. (44) who provide a software package for performing method 2 optimisations. A tool by Teed and Silva (45) similarly implements a Method 2, but also varies model structure when searching. While some authors prefer manually adjusting model parameters, several optimisation algorithms have also been applied to Method 2, including simulated annealing (45), a Davidon-Fletcher-Powell optimiser (46), subspace trust region methods (40), and the Nelder-Mead downhill simplex method (40, 43).

Instead of fitting to processed ‘summary’ data (time constants and steady states), we can also fit simulated currents directly to measured currents from the same protocols: in this manuscript we term this ‘*Method 3*’. Like Method 2, Method 3 is applicable to Markov style models. As Method 3 does not require calculation of time constants and steady states it is insensitive to errors in this process and can be more computationally efficient. A downside is that full current traces are more difficult to obtain, and require the experimenter to have published their findings in digital format, rather than printed tables or summary graphs.

The applicability of Method 3 to Markov models was exploited by Balser et al. (47), who used numerical optimisation and whole-current fitting to find the parameters of a model describing a cardiac potassium current. Similarly, Irvine et al. (48) formulated a Markov model of the cardiac fast sodium current and fit it using Method 3 (although they used additional data sources beyond whole-cell currents). Several algorithms have been used for the optimisation step in Method 3, including the Levenberg–Marquardt algorithm (47, 49), Nelder-Mead (47), simulated annealing (48), differential evolution (50), and genetic algorithms (51, 52), as well as hybrid methods combining a multiple algorithms (53).

A number of studies have been published which investigate the identifiability of Hodgkin-Huxley models from current recordings. Willms et al. (29) highlighted advantages of Method 3 over Method 1, and provided a software tool for Method 3 fitting (54), while an extensive analysis for *voltage-independent* channels was given by Milescu et al. (55). Csercsik et al. (56) investigated a procedure falling somewhere between Methods 1 and 3, where the parameters underlying voltage-dependence of steady states were determined, but time constants were fit separately for different voltages. Walch and Eisenberg (57) considered the problem of estimating both time constants and steady states independently for several voltages, and concluded that in this case only the time constants were identifiable. As Csercsik et al. (56) point out both these analyses do not fully benefit from the voltage-dependence implied by the equations, which again points to possible benefits for a Method 3 approach, where the full model is fit to currents recorded at several voltages.

Method 3 fit to current traces directly, without calculation of time constants or steady states, yet used protocols designed specifically for this purpose. A next natural step then, is to reconsider the established protocols and to ask what type of protocol would work best for Method 3. We define ‘*Method 4*’ as the process of fitting directly to currents measured using a single information-rich optimised protocol.

Perhaps the first demonstration of Method 4 was by Millonas and Hanck (58), who fit a Markov model of a cardiac sodium current using a protocol consisting of random fluctuations between two voltages (‘dichotomous noise’). This method was later applied to design protocols specifically to differentiate between competing models (59–61). A brief mention also occurs in Gurkiewicz and Korngreen (51), which discusses the benefits of Method 3 for semiautomatic ‘high-throughput’ construction of ionic current models and notes that it could be applied to ‘non-standard protocols’. Fink and Noble (62) used identifiability analysis and simulation to show that standard voltage protocols can be considerably shortened without losing information. Importantly, they showed that protocols can be created that provide information on parameters for multiple cardiac currents: it may be possible to formulate protocols without extensive pre-knowledge of the system (as is typical for standard protocols), and that protocols might be designed which are robust to changes induced by e.g. drugs or mutations. Subsequent work by Clerx et al. (63) showed that the identifiability analysis employed by Fink and Noble (62) could be extended to take into account experimental errors. Finally, a paper by Beattie et al. (14) proposed and tested a protocol design based on sinusoidal voltage clamps. Their work showed that a model could reliably be fit to the resulting current measurements, and that models created this way outperformed published models in predicting the response to conventional voltage clamp protocols and an AP-waveform protocol.

Having discussed the four methods, we should point out that many papers do not stick to using a single method, but use different approaches for different model parameters (64). In the remainder of this paper, we formulate the four methods that characterise the approaches discussed above, and evaluate their performance on a previously-published experimental data set.

## MATERIALS AND METHODS

### Current model

To model the hERG current, we use a two-gate Hodgkin-Huxley model found to provide excellent predictions by Beattie et al. (14) (described there in equivalent Markov model form). The current is defined by

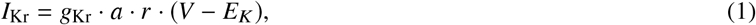

where *g*_Kr_ is the maximum conductance, *a* and *r* are gating variables (defined below), *V* is the trans-membrane potential and *E_K_* is the reversal potential for potassium ions (note that, although the measurements we used were from CHO cells expressing hERG1a, we use the shorthand terms *I*_Kr_ and *g*_Kr_ throughout this paper). The Nernst equation was used to calculate a separate *E_K_* value for each cell:

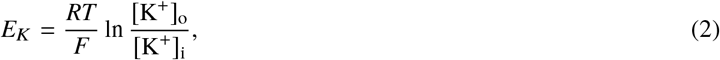

where *R* is the gas constant, *F* is the Faraday constant, [K^+^]_o_ and [K^+^]_i_ are the bath (outside membrane) and pipette (inside membrane) concentrations of potassium ions and *T* is the cell-specific temperature measured by Beattie et al. (14). In the 9 cells used in this study, calculated values for *E_K_* ranged from −88.45mV to −88.30mV with the precise value depending on the temperature at the time of recording.

The two processes represented independently by this Hodgkin-Huxley model are *activation* (*a*) and *recovery* (*r*). Both variables *a* and *r* are dimensionless and in the range [0, 1]. Increases in *a* correspond to ‘activation’ and decreases in *a* to ‘deactivation’; while decreases in *r* correspond to ‘inactivation’ and increases in *r* to ‘recovery’ [from inactivation].^1^ We can represent the model as two independent gating reactions

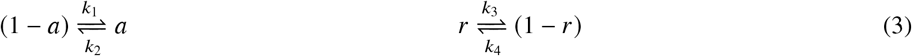

where *a* is the fraction of channels in the activated state, (1 − *a*) is the fraction in the deactivated state, *r* is the recovered fraction, (1 − *r*) the inactivated fraction, and *k*_1_ to *k*_4_ are the voltage-dependent transition rates. The ordinary differential equations governing *a* and *r* are then derived with mass-action kinetics and can be written in the form:

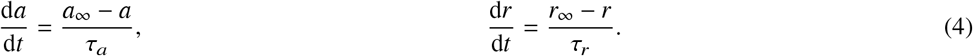

Where *a*_∞_ and *r*_∞_ denote voltage-dependent steady-states and *τ_a_* and *τ_r_* denote voltage-dependent time constants defined in terms of the transition rates as

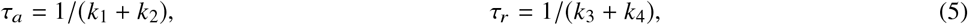

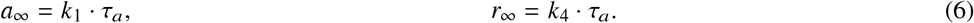

The voltage dependencies of the transition rates are defined using an Eyring-derived exponential formulation as

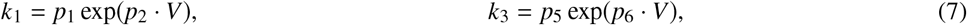

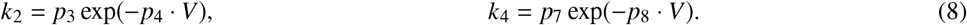

The model parameters to be inferred are therefore the kinetic parameters *p*_1_, *p*_2_, …, *p*_8_ and the conductance *p*_9_ = *g*_Kr_. All model parameters are constrained to be strictly positive: *p_i_* > 0 for *i* ∈ 1, 2, …, 9.

### Experimental methods

The experimental data used in this study are taken from Beattie et al. (14). In short, manual patch-clamp recordings were performed at room temperature (between 21–22°C) in CHO cells stably expressing hERG1a (which encodes the alpha subunit of the channel carrying *I*_Kr_). Recordings were taken from nine cells, and seven protocols in each cell (we will refer to these as Cells #1 to #9, and Pr1 to Pr7, with the numbering matching the original publication). After the final protocol was completed, the *I*_Kr_ blocker dofetilide was washed in, and all protocols were repeated. Each cell’s data was then post-processed by first leak-correcting the signals recorded both in the presence and absence of dofetilide, before subtracting the *I*_Kr_-blocked signal from the unblocked one to remove any endogenous currents. For this study, we used the already leak-corrected and dofetilide-subtracted data as published on https://github.com/mirams/sine-wave. The first protocol, Pr1, did not elicit strong currents in any of the cells, and so was not used in this study.

Following Beattie et al. (14), capacitance artefacts were removed from the experimental data by discarding the first 5ms after each discontinuous voltage step. To obtain similar results from simulated protocols, the same filtering was applied to all simulated data.

The six protocols used in this study are shown in Figure 1. The first four, Pr2–5, are adaptations of common step protocols used to characterise *I*_Kr_. Specifically, Pr2 is used to estimate a single time constant of activation (for *V* = +40mV), Pr3 is used to estimate the steady state of activation, Pr4 is used to estimate time constants of inactivation, and Pr5 provides data about both time constants, as well as the steady state of inactivation. Pr7 is a novel sinusoidal protocol introduced by Beattie et al. (14), and is intended to provide the same information in a much shorter time. It consists of an initial step to +40 mV, designed to trigger a large current, followed by a section consisting of the sum of three sine waves. Finally, Pr6 is a collection of AP wave forms, representing the membrane potential under physiological and pathological conditions. As in Beattie et al. (14), we used Pr6 as a *validation* protocol, while either Pr7 or the set Pr2–5 were used for fitting. Note that the full set of protocols was run on every cell.

**Figure 1:**
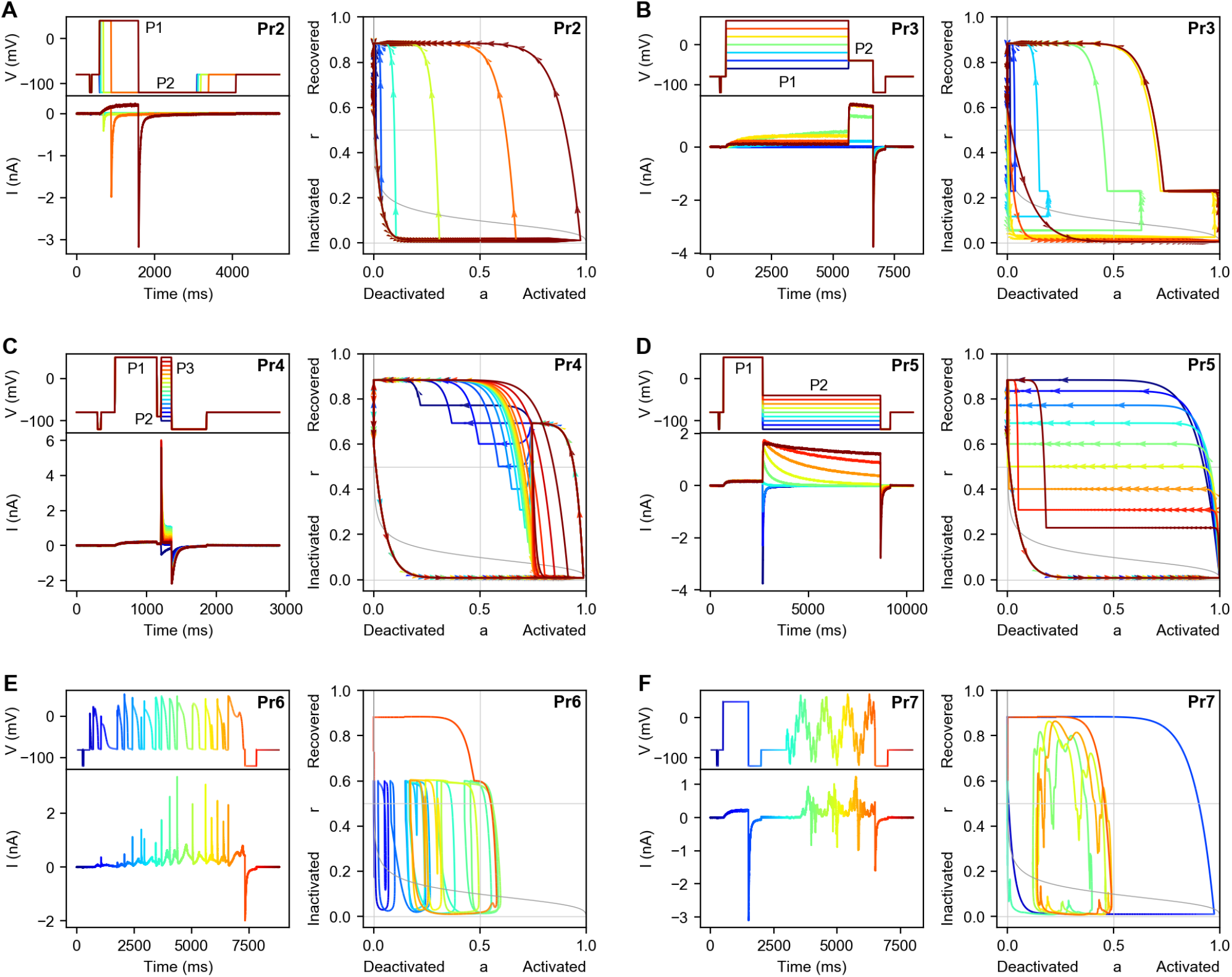
Voltage protocols, currents measured in Cell #5, and simulated phase diagrams for all 6 protocols (Pr2–7) used in this study. Simulations for the phase diagrams were performed with the Cell #5 parameters obtained in Beattie et al. (14). (*A*) Pr2 is used to measure a time constant of activation. We show the voltage step protocol (*Top left*), the resulting current as measured in Cell #5 (*Lower left*), and a phase plane diagram (*Right*). The protocol is repeated 6 times, with an increasing duration of the P1 step for each repeat. This is indicated in the plots by using a different colour for each repeat. (*B*) Pr3 is used to approximate the steady state activation curve. It is repeated 7 times, with a different voltage for the P1 step at each repeat. (*C*) Pr4 is used to measure time constants of inactivation, it repeats 16 times with a different voltage for step P3. (*D*) Pr5 is used to measure time constants of activation and inactivation, as well as steady state inactivation. It has 9 repeats with a different voltage for step P2. (*E*) Pr6 consists of several (healthy and pathological) AP wave forms, and was used as an independent validation data set in this study. The colours in this plot indicate time, so that the voltage and current traces can be related to the phase plane trajectory on the right. (*F*) Pr7 is the sine wave protocol introduced by Beattie et al. (14). As in panel E, colour is used to indicate time information.

A detailed description of all protocols and the associated analysis methods is given in Supporting Material S1. In analysing these protocols, we found it useful to perform simulations (using the parameters obtained by (14)) and plot the model state in a two-dimensional *phase diagram* as shown in Figure 1.A guide to using these diagrams for analysis is provided in Supporting Material S1.1.

The voltage step protocols Pr2–5 can be used to derive a set of graphs which characterise the current, shown in Figure 2 for all nine cells used in this study. They are commonly referred to as the ‘steady of activation’ (Figure 2.A), ‘steady state of inactivation’ (Figure 2.B), ‘[Peak/tail] IV curve’ (Figure 2.C), ‘time constant of activation’ (Figure 2.D), and ‘time constant of inactivation’ (Figure 2.E). In the remainder of the paper, we will refer to the curves in Figure 2 as the *summary curves*. Such curves have been considered historically as imperfect direct approximations of the *a*_∞_, *r*_∞_, *τ_a_*, and *τ_r_* variables within a model, or simply as relevant summaries of the electrophysiology produced by following a fixed experimental procedure. To distinguish the summary curves from the model variables, we will use a ‘tilde’-notation to denote the experimental/simulated summary curve values: 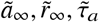 and 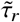.

**Figure 2:**
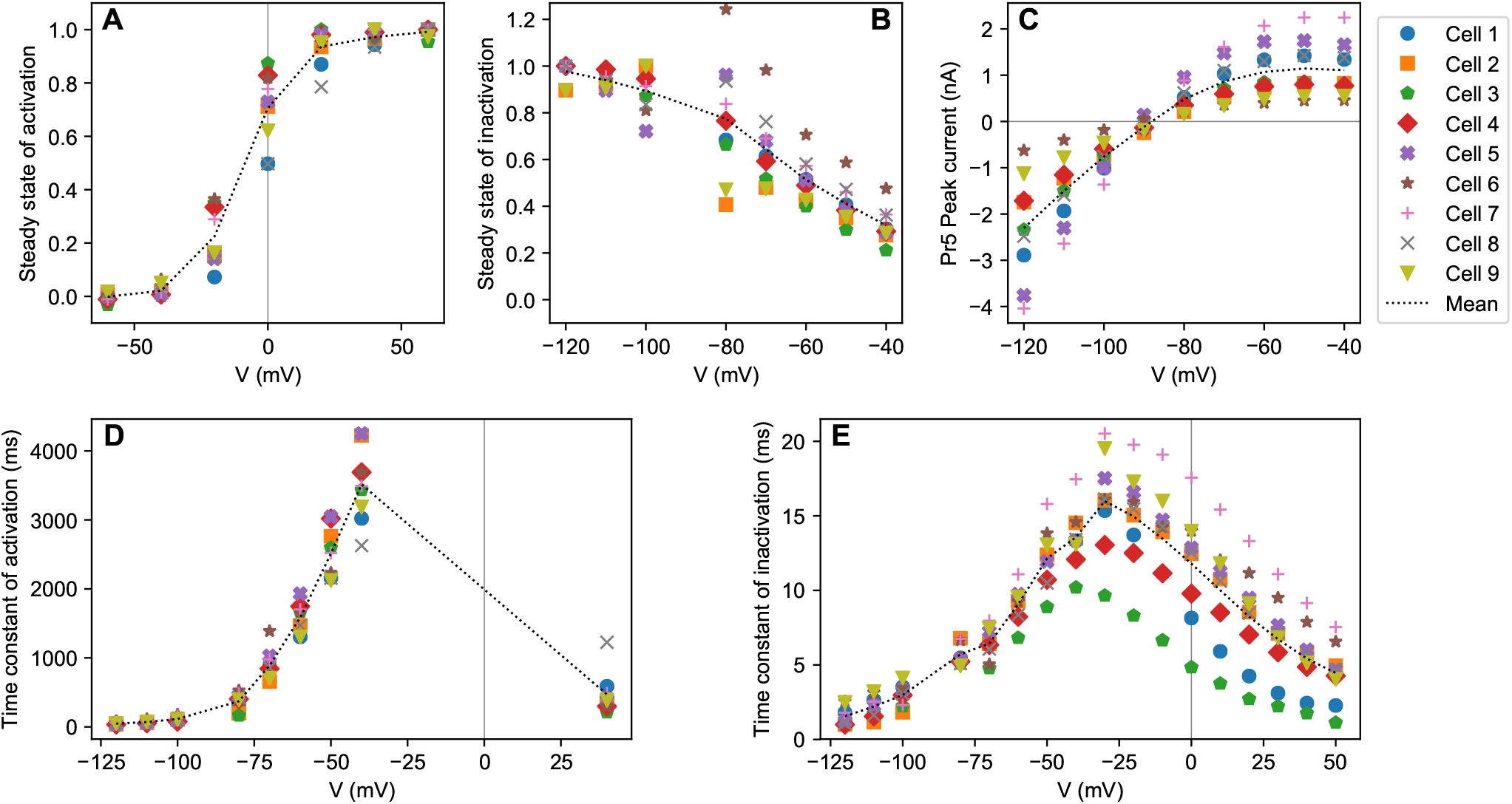
Summary curves calculated from the voltage-step protocols Pr2–5 for all nine cells. The mean for all cells is indicated with a dotted line. Note how the general trend, not always obvious from the single-cell results, is clearly captured by the line connecting the means. (*A*) The activation curve, which approximates *a*_∞_, derived from Pr3. (*B*) The steady-state of inactivation, which approximates *r*_∞_, derived from Pr5. (*C*) The IV curve (peak current during the P2 step of Pr5) plotted against the P2 voltage. (*D*) The estimated time constant of activation (approximating *τ_a_*). Values for *V* < 0mV were estimated from Pr5, the single value at 40mV from Pr2. (*E*) The estimated time constant of inactivation (approximating *τ_r_*). Values for *V* < −30mV were estimated from Pr5, values for −40mV and upwards from Pr4.

### Four ways of fitting

#### Method 1: fitting model equations directly to summary curves

We now describe the four methods of fitting in detail. In Method 1, we write out equations for *a*_∞_, *r*_∞_, *τ_a_*, and *τ_r_* in terms of the parameters *p*_1_, *p*_2_, …, *p*_8_, and then try to fit these directly to the summary curves 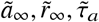, and 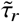. The procedure for doing so closely follows that of Hodgkin & Huxley, and in fact the procedure for finding *p*_1_ to *p*_8_ can be done entirely with pen and paper, although we will use a simulation to estimate the 9th parameter *g*_Kr_. First, we focus on the steady state of activation (Eq. 6) and substitute in the parameters *p*_1_, to *p*_4_:

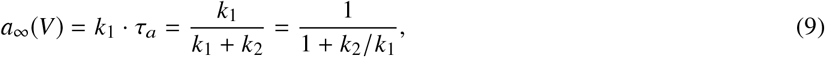

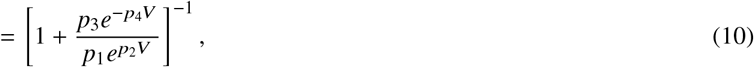

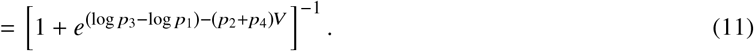

This can be rewritten as a ‘Boltzmann curve’

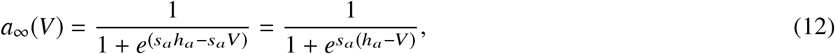

where *h_a_* is the ‘midpoint of activation’ (the point where *a*_∞_(*V* = *h_a_*) = 0.5), and *s_a_* is the slope of the activation curve at *V* = *h_a_*. Assuming 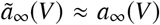, values for *h_a_* and *s_a_* can then be obtained numerically (optimising on sum of square error) or by plotting *a*_∞_ versus *V* and reading the values from the graph. At this stage we have an equation for *a*_∞_(*V*), and two constraints on the parameters *p*_1_, *p*_2_, *p*_3_, *p*_4_, namely:

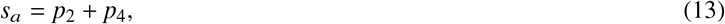

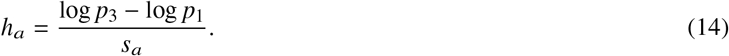

Next, we rewrite Eq. 6 to find

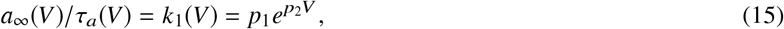

which allows us to infer values for *p*_1_ and *p*_2_ by plotting the quantity 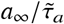 versus *V* (using the fitted equation for *a*_∞_ rather than the measured values to reduce noise) and fitting an exponential curve. Note that since *a*_∞_ ≈ 0 for values below −60 mV, any data points for low voltages will contribute very little to the final fit. As a simple alternative, we can plot the quantity 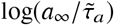 and fit with a linear equation log(*p*_1_) + *p*_2_*V* (note that this is equivalent to doing a graphical fit using semi-logarithmic graph paper). This provides us with more reliable values for *p*_1_ and *p*_2_, after which we can use *p*_4_ = *s_a_* − *p*_2_ and *p*_3_ = *p*_1_*e^s_a_h_a_^* to find the remaining activation parameters.

Using a similar procedure (but with a slight change in signs), we plot the logarithm of 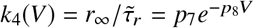, fit to find *p*_7_ and *p*_8_, and then use *p*_6_ = −*s_r_* − *p*_8_ and *p*_5_ = *p*_7_*e^s_r_h_r_^* to find all four inactivation parameters.

Finally, we find a value for the conductance parameter *g*_Kr_ by performing a simulation of Pr5, deriving an IV-curve, and then calculating the scaling factor that minimises the sum-of-squares error between the simulated and experimental IV curves.

#### Quantifying goodness-of-fit

To evaluate the goodness-of-fit from Method 1, we derive an error function and evaluate it with the obtained parameter values. We write ***θ*** = {*p*_1_, *p*_2_,…,*p*_9_} for the parameters and introduce symbols representing the experimental and measured data sets. To denote a cell’s set of experimentally approximated midpoints of activation we use 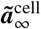, while 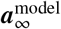 are used to indicate the value of the model variable (*a*_∞_), evaluated at the same voltages. Similar notation is used for the midpoint of inactivation and both time constants. Next, we define the root mean squared error (RMSE) between two data sets ***x*** and ***y*** as

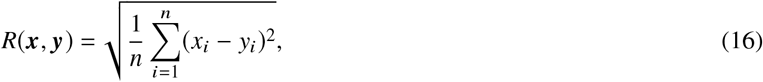

where both data sets have equal length *n*. Using this notation, we can write the RMSE between experimental results 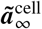 and model values 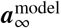 as 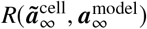. With this notation, we can now define the error criterion as a weighted sum of RMSEs:

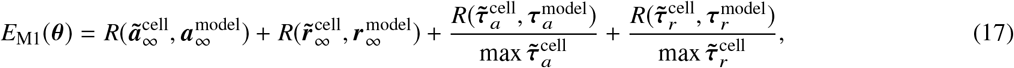

where *n*_*a*_∞__ = 7, *n*_*r*_∞__ = 8, *n*_*τ*_*a*__ = 9, and *n*_*τ*_*r*__ = 17. Note that this is a cell-specific measure, as both time constants are normalised with respect to the largest value found in the used cell data. This weighting corrects for differences in the scaling of the four RMSEs, ensuring that none of them dominate in the end result. No normalisation is needed for the steady-states, which are already constrained to the range (0, 1). In contrast to measures introduced below, *E*_M1_ is invariant with respect to conductance (*p*_9_). Finally, we note that an alternative Method 1 could be created by using numerical optimisation to minimise *E*_M1_, this is explored further in Supporting Material S3.6, it did not create a more predictive model than the method presented here.

#### Method 2: fitting simulated summary curves

In Method 2, we accept that the summary curves are imperfect approximations of the model variables, and so we base our fitting on *simulated experiments* of Pr2–5, analysed using the same methods as for the experimental data to arrive at simulated versions of the summary statistic curves. This gives us two sets of summary curves, one simulated and one experimental, using which we can define an *error measure* that quantifies the goodness-of-fit. By varying the parameters and repeating the simulations we can then find a set of parameters that minimises this error.

To formulate the error measure, we again write 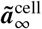 to denote a cell’s set of experimentally approximated midpoints of activation, and we introduce the notation 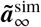 for its simulated counterpart. In this measure, we use the IV-curve rather than the steady-state of inactivation as (1) it contains the same information (albeit with a different scaling); (2) unlike 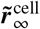, it does not suffer from numerical issues near *V* = *E_K_* (see Supporting Material S1.5.1); and (3) it contains information about *g*_Kr_, which is lacking from the other summary curves. Similar symbols denote the remaining summary curves: midpoint of inactivation 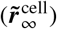, time constant of activation 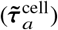, time constant of inactivation 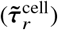 and the IV-curve from Pr5 (**IV**^cell^).

The error to minimise for each cell is defined as a weighted sum of RMSEs

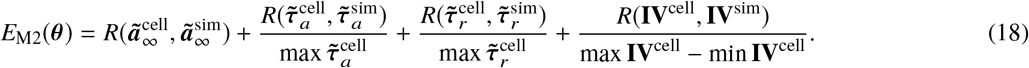

The number of data points was the same for each cell, with *n_τ_a__* = 9, *n_τ_r__* = 17, *n_a_∞__* = 7, and *n_IV_* = 9 (for a total of 42 cell-specific data points). As in *E*_M1_, weighting is used here to give every term equal influence.

#### Method 3: fitting current traces from traditional protocols

In Method 3, we forgo the summary curve calculation altogether and simply perform ‘whole trace fitting’ on the currents resulting from the ‘traditional protocols’ Pr2–5. Writing 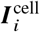 for the current recorded from protocol *i*, and 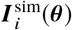 for the simulated current in protocol *i* with parameters ***θ***, we define the function to be optimised as a normalised RMSE:

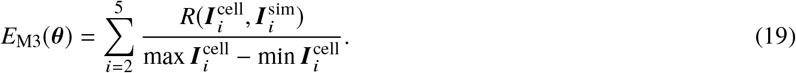

Note that the weighting here is not strictly necessary for the optimisation procedure, but is used to enable *E*_M3_ value comparisons between cells.

#### Method 4: fitting current traces from an optimised protocol

In Method 4, we define a similar normalised RMSE measure based on fitting only the current under the sinusoidal protocol, Pr7:

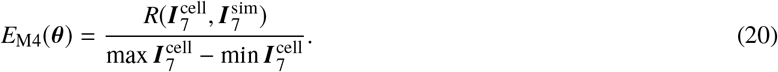

As in *E*_*M*3_, the weighting used here allows us to compare values between cells.

### Validation and cross-comparison

To compare the results of the four fitting methods, we applied each method to each cell, resulting in 4 parameter sets per cell. Next, we simulated the AP waveform protocol (Pr6) with all parameter sets, and compared these to the corresponding measurements. Note that Pr6 was not used in any of the fitting methods, so that this constitutes an independent validation. The results were quantified using a normalised RMSE:

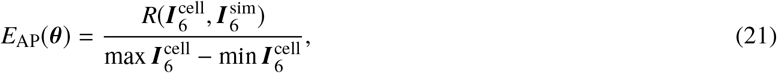

where *R* is defined by Eq. 16, as before.

In addition to the independent validation, we performed cross-validation by testing how well models fit with one method could predict the fitting data used by the others. This was shown visually, and was quantified by evaluating *E*_M1_–*E*_M4_ on the best result found for each cell/method.

Multi-cell measures of the fitting methods’ performance were defined as the mean average of the error measure over all nine cells. Writing *E*_AP,k_ for the RMSE in cell k on the AP signal, we defined

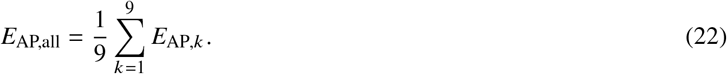

Note that *E*_AP,*k*_ already contains a term to normalise the error with respect to the magnitude of the current in cell *k*, so that no further weighting is required. Combined error measures for *E*_M1_ through to *E*_M4_ were defined in the same manner.

### Minimising the error measures

Methods 2, 3, and 4 all proceed by finding a parameter set that minimises an error function. In previous work we found that the global optimisation algorithm CMA-ES (65) provided good fits for a range of models from the ion channel to cell scale (14, 66–69), and was both fast and behaved reproducibly (returning the same answer when started from different initial guesses in the parameter space). For Methods 2, 3, and 4 we used a CMA-ES population size of 10, and halted the optimisation only when the objective function changed by less than 10^−11^ per iteration for 200 successive iterations. Other important aspects of making our optimisation strategy reliable were (i) placing physiologically-inspired bounds on the parameter space; (ii) searching in a log-transformed space; (iii) reducing solver tolerances to eliminate numerical noise in simulation output; and (iv) testing reliability by running repeated fits from different starting points.

Constraints on the parameter space were set as in Beattie et al. (14), and included upper and lower bound for the parameters *p*_1_ to *p*_9_ but also restricted the value of the reaction rates *k*_1_ to *k*_4_. The resulting boundaries are visualised in Figure 3.A and B. Details are given in Supporting Material S2.2.

**Figure 3:**
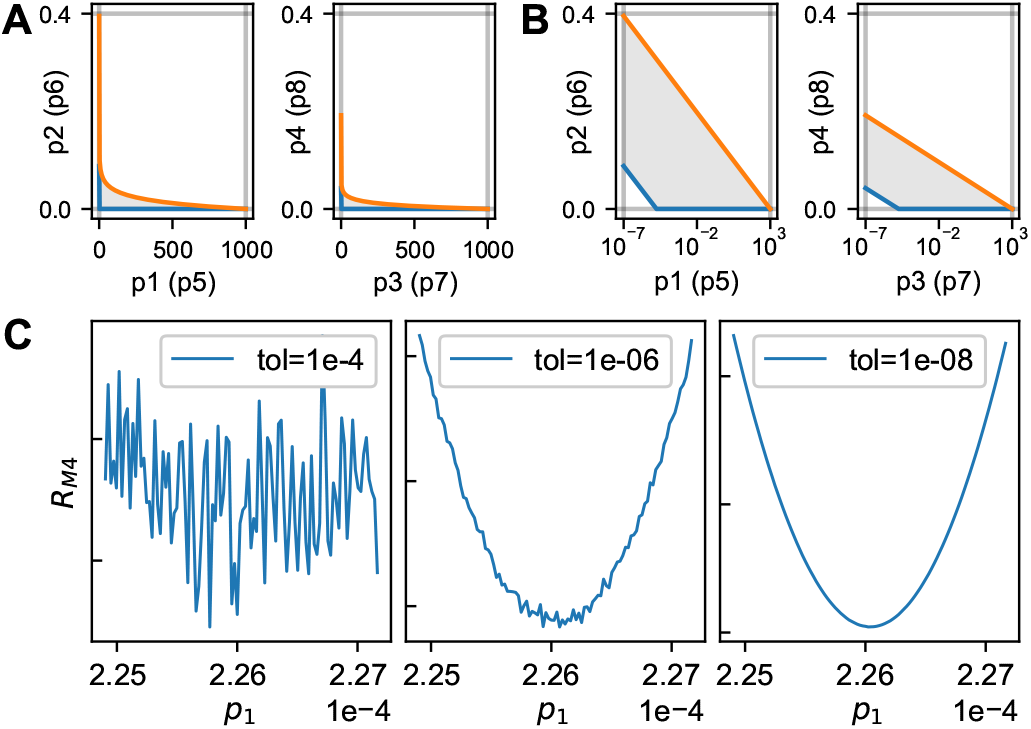
Boundaries and transforms on the parameter space and the effect of solver tolerances. (*A*) Constraining the transition rates *k*_1_–*k*_4_ (over physiologically-relevant voltages) to relevant timescales creates a 2-dimensional boundary on each parameter pair (Supporting Material S2.2). The grey region indicates valid samples, with an upper rate constraint in orange and a lower rate constraint in blue. (*B*) the same parameter constraints with a log transform on one of the parameters. Note this increases the size of the feasible region for off-the-shelf optimisers that know nothing about the problem, and we later see the objective function becomes reasonably convex and symmetric under this transform (see Figure 11). (*C*) Using default tolerances of adaptive solvers can lead to numerical noise in the function being optimised. This can be remedied by selecting lower tolerances.

Several studies (70–72) have found that, for strictly positive parameters (i.e. for all 9 parameters in our model) optimisation performance can be improved searching in a log-transformed parameter space. In at least some cases, this has been shown to turn non-convex (i.e. hard) problems into simpler convex ones (73). For Methods 2, 3, and 4, we used a log-transform on parameters *p*_1_,*p*_3_,*p*_5_, and *p*_7_.

We used an adaptive step time solver for simulations of Pr6 and Pr7. As shown in Johnstone (74), simulating with lax tolerance settings can lead to (seemingly random) fluctuations in the error measure for nearby values of ***θ***, which has the effect of creating several local minima in otherwise smooth regions. To combat this, we set absolute and relative error tolerances of 10^−8^. This is shown in Figure 3.C. For the voltage-step protocols Pr2–5, we circumvented the issue entirely by using the model’s analytical solution for fixed voltages to calculate the time course at each step.

When running the optimisations for Methods 2, 3, and 4, each fit was run several times from different starting points, sampled uniformly from within the transformed space shown in Figure 3.B. Fifty repeats were run per cell for every method, and the best result (lowest error) was used as the final fit for that cell/method. We comment on the reliability of these fits below.

### Software and algorithms

Simulations were performed in Myokit (75), using CVODE (76) for Pr6 and Pr7 simulations with tolerances as described above, and an analytical solver for Pr2–5. Further analysis was performed in Python 3.6 using NumPy/SciPy (77). When deriving time constants, fits to exponential curves were performed using the Nelder-Mead downhill simplex algorithm implemented in SciPy. All other fits were performed using CMA-ES (65), via the PINTS (78) inference framework. When calculating benchmarks, Methods 2, 3, and 4 were run on a machine with 24 Intel Xeon 2.2GHz CPU cores (48 with hyperthreading), with no more than 4 optimisations running at the same time. All data, code, results, and figures are available to download from https://github.com/CardiacModelling/FourWaysOfFitting.

## RESULTS

We now discuss the results of fitting a model with each of the four methods, using Cell #5 as an example in our figures. Figures for all nine cells can be found online at https://github.com/CardiacModelling/FourWaysOfFitting. For each fit, we discuss the quality of fit, but also investigate whether models made with the other methods do well at predicting the methods’ fitting data.

In Method 1 we fit model equations (e.g. *a*_∞_) to experimental approximations (e.g. 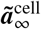). Figure 4 shows the experimentally derived summary curves as black squares, while the blue line (‘Fit 1’) represents the quality of fit obtained by Method 1. Note that these lines are plotted directly from Eqs. 5 & 6, and do not involve simulation. Similar lines are shown for models fit with Methods 2–4, labelled as ‘predictions’ in the figure. As Figure 4 shows, the lines drawn from the Method 1 model fit the data well, while the models fit through simulation (Methods 2–4) show a notable mismatch for both steady states and time constants. However, as the summary curves approximate but do not equal the model variables, a good fit in this figure is not necessarily desirable. This is discussed further in Supporting Material S1.6.

**Figure 4:**
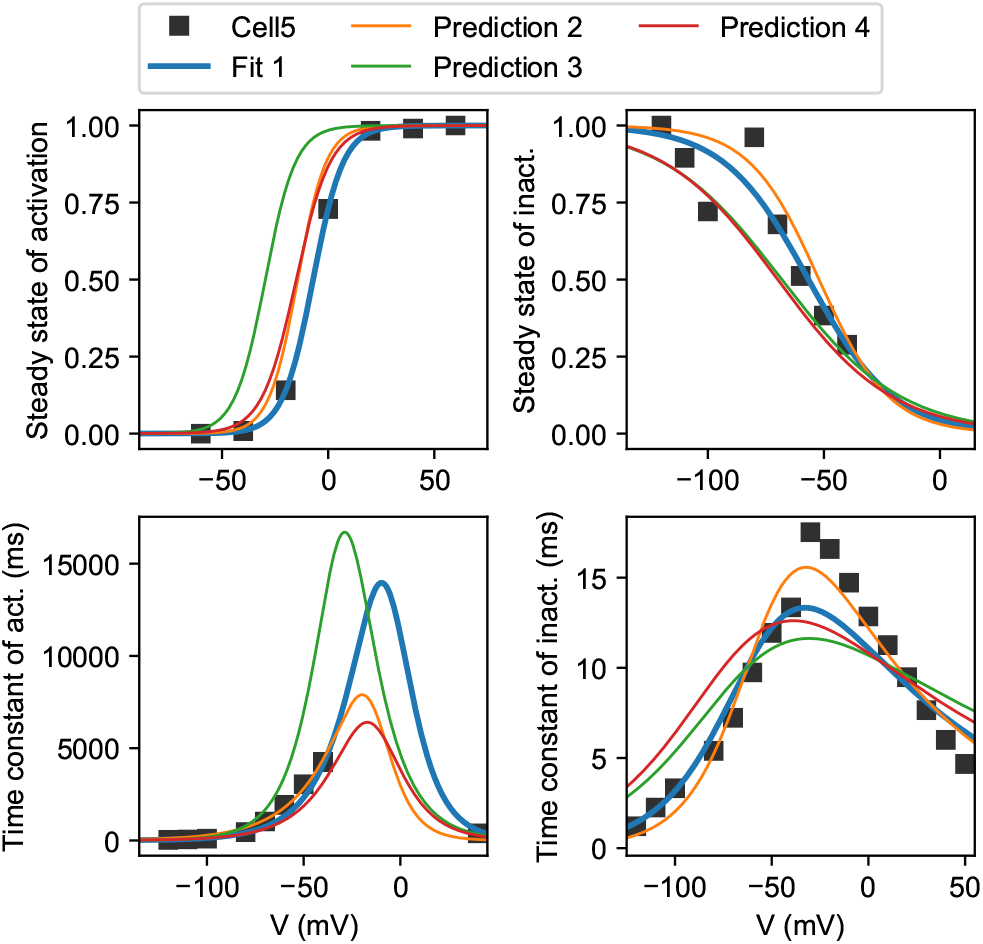
Method 1 goodness-of-fit and cross validation on Cell #5. Experimental approximations of the steady states (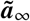 and 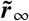) and time constants (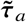 and 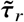) are shown, derived from measurements in Cell #5. The model curves for *a*_∞_(*V*), *r*_∞_(*V*), *τ_a_*(*V*) and *τ*(*V*) are shown for models fit with each of the four methods. Note that only Method 1 was trained on this data — making this a goodness-of-fit figure for Method 1, while for the other methods this figure shows a prediction.

In Figure 5, we again plot the experimentally derived data points (e.g. 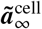), but instead of comparing them to model variables, we compare them to simulation results (e.g. 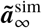). Method 2 attempts to minimise the discrepancy between the two, and achieves this well (‘Fit 2’ in Figure 5). By contrast, the predictions from a model made with Method 1 now show a clear shift in the steady state of activation curve. Interestingly, only the method-2 derived model shows a close fit to the time constant data, although none of the methods are able to match the steep peak in the 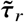 points well, which may be due to the fact that points left of the peak were derived from Pr5, while the right-most points are obtained from Pr4.

**Figure 5:**
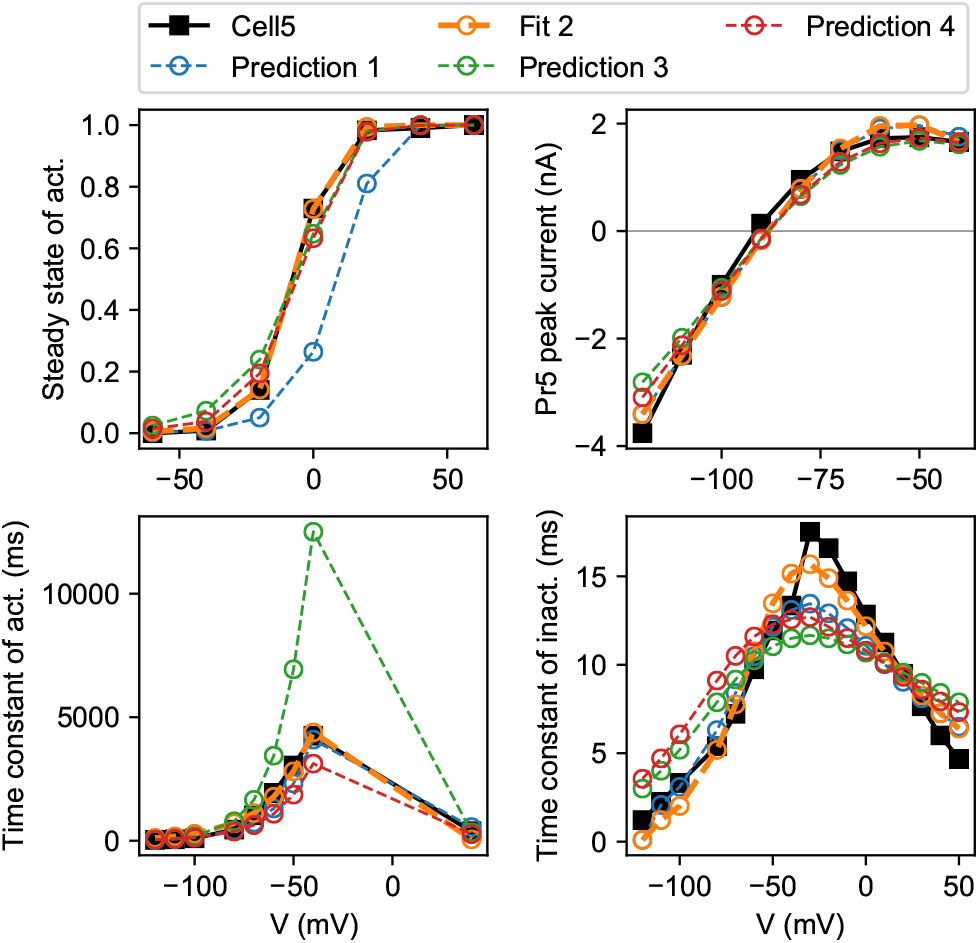
Method 2 goodness-of-fit and cross validation on Cell #5. Experimental approximations of the steady state of activation, the IV curve, and both time constants are shown. By simulating the protocols and performing the same analysis, similar approximate points were derived for each of the four methods. The quality of fit for Method 2 is shown, along with the predictions from models made with Methods 1, 3, and 4.

Figure 6 shows selected portions of the currents elicited by Pr2–5. Method 3 minimises the discrepancy between simulated and measured currents, and provides a good fit (‘Fit 3’ in the figure) although some differences can still be seen. Models made with Methods 1 and 2 do not generally predict the observed currents well, although the qualitative behaviour is correct in all cases. Models with Method 4 provide better predictions here, although interestingly, only Method 3 gets the slope of the Pr3 currents during the P2 step right, indicating a difference in the deactivation properties of Method 3 parameters compared to the others. The negative currents in Pr4 appear challenging for all methods, while Method 3 and 4 fits match well on the higher potential steps.

**Figure 6:**
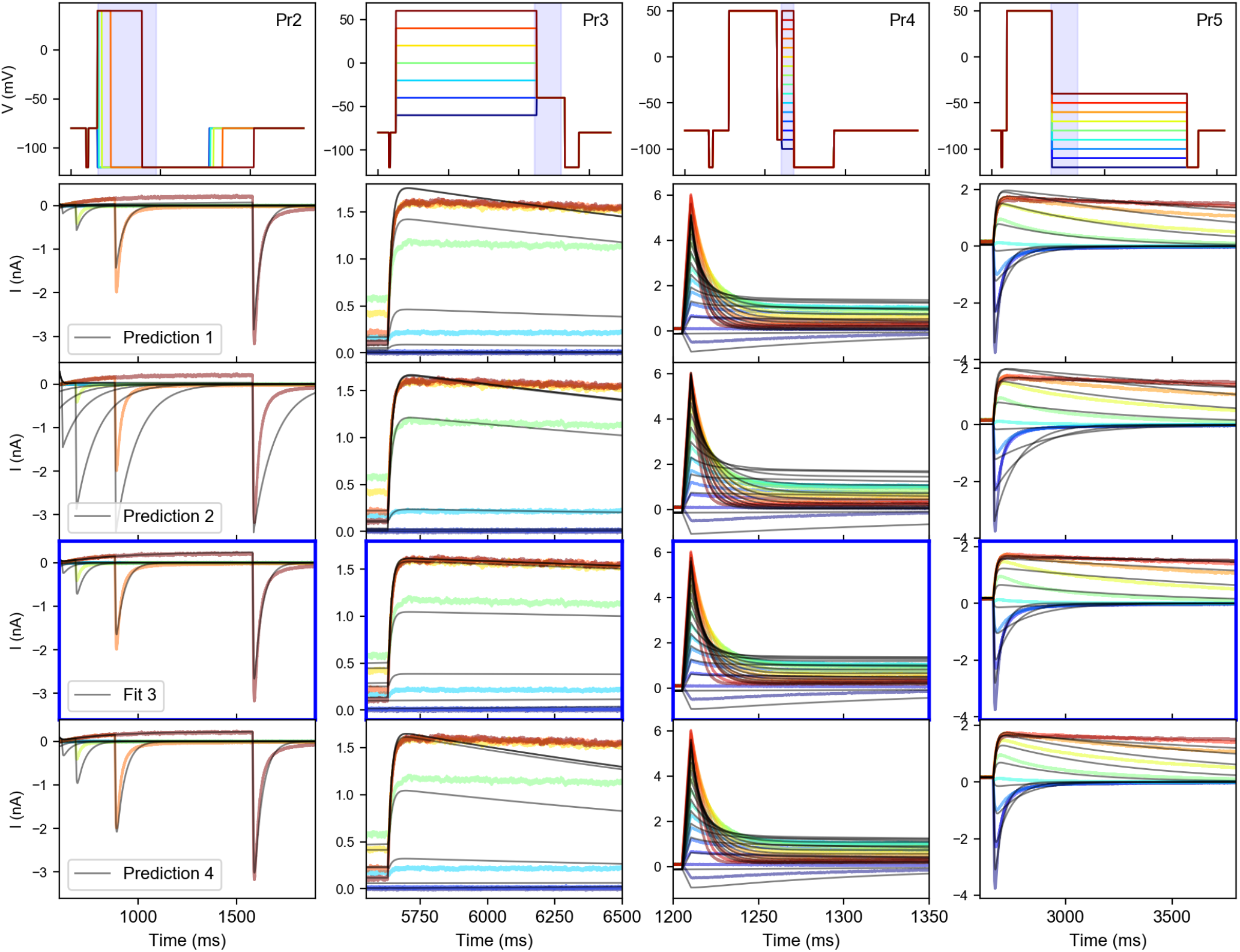
Method 3 goodness-of-fit and cross validation on Cell #5. The top row shows the voltage protocols Pr2–5, with different colours used for each sweep. The same colours are used in each of the panels below to show the corresponding experimental data, while simulation results are shown in black. In the top row, the full time-span of each protocol is shown, but the plots shown below each zoom in on a specific region, indicated by the shaded area in the top row. Quality-of-fit is highlighted with blue borders for Method 3 (‘Fit 3’), alongside predictions from models made with Methods 1, 2, and 4.

Next, we inspected the capability of the different models to predict the currents from the sine wave protocol (Pr7), as shown in Figure 7. The Method 4 model obtained a good fit to the data, although some differences can be seen in the deactivating part of the initial voltage step. Predictions from the model made with Method 3 were relatively good, while models fit with Method 1 and in particularly Method 2 performed poorly at predicting Pr7 currents.

**Figure 7:**
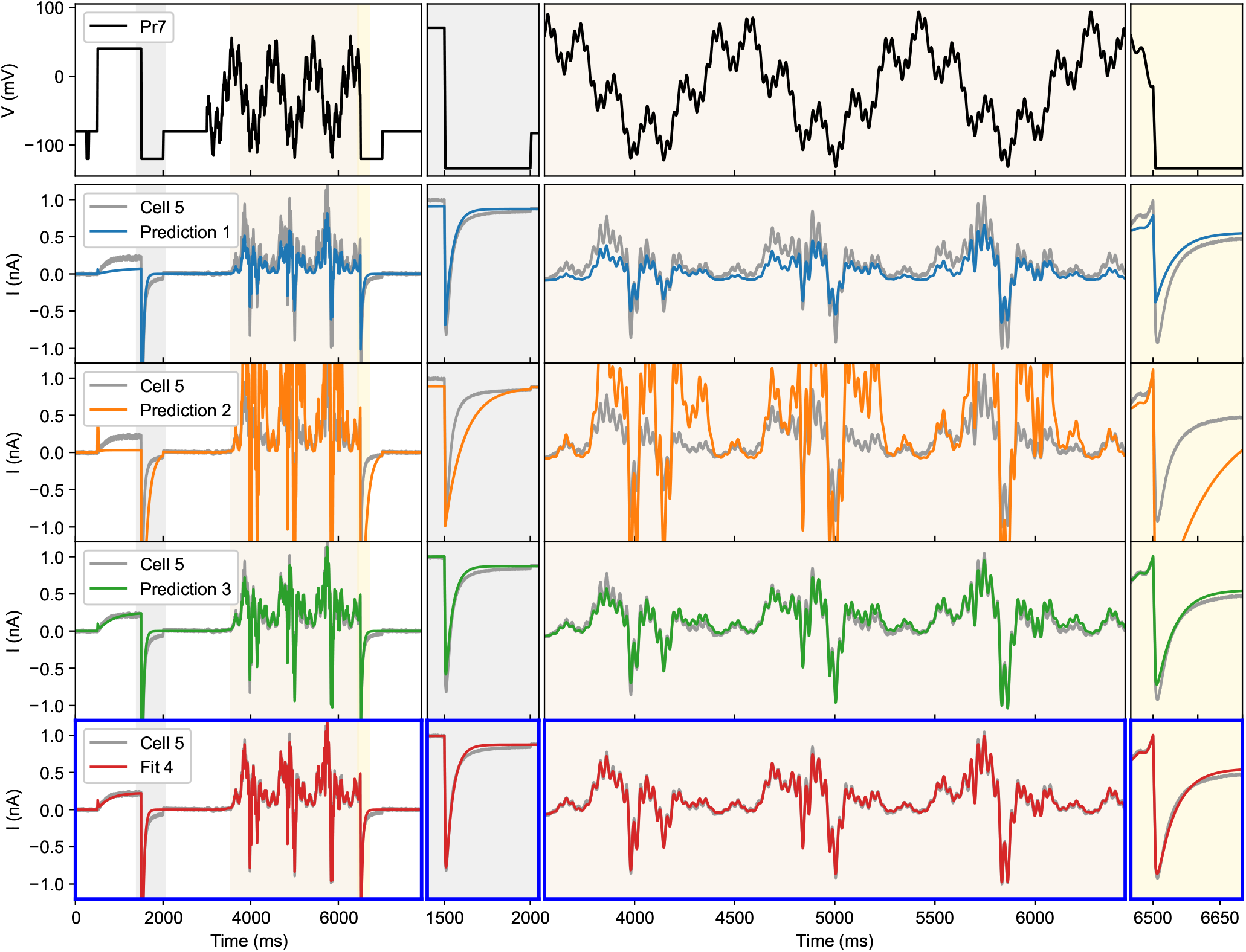
Method 4 goodness-of-fit and cross validation on Cell #5. The top panel shows the voltage protocol (Pr7), while the three panels below show the same current recording (grey line) and each row shows a prediction from a model made with Methods 1, 2, and 3 respectively (coloured lines). The bottom panel, highlighted with blue borders, shows the fit for Method 4 (‘Fit 4’).

Figures 3–6 showed that all four methods achieved good fits—judged by their own criteria — while the quality of predictions outside of the fitting data varied. The AP waveform is an attempt to test predictions for the most physiologically-relevant *I*_Kr_ behaviour. A visual comparison of predictions made with models from each of the four methods is shown in Figure 8, again with the Cell #5 results used as an example. In this cell, the predictions from Method 2 were generally poor, while Methods 3 and 4 provide much closer fits.

**Figure 8:**
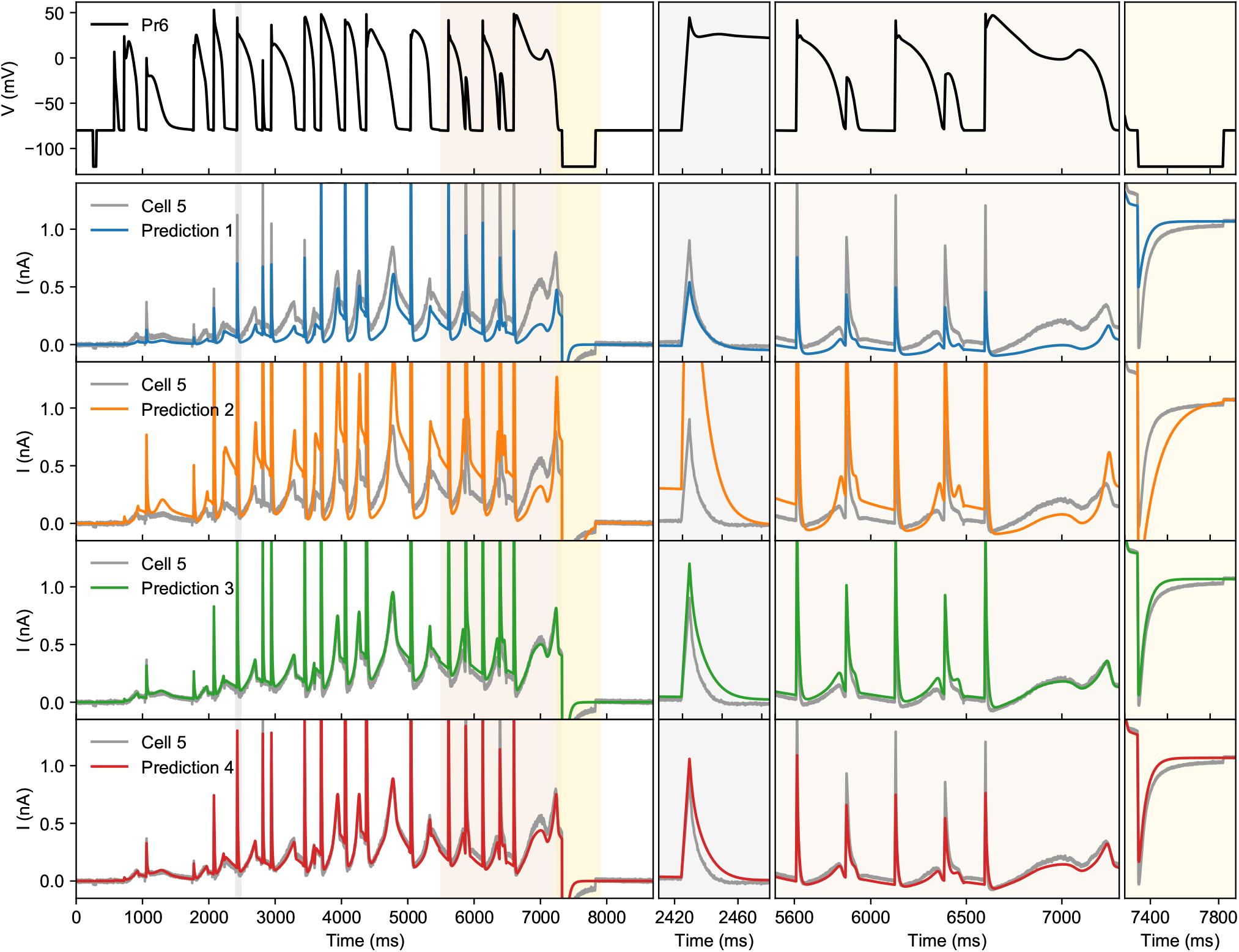
Validation on the AP-waveform signal (Pr6). The voltage protocol is shown on the top row, with subsequent rows coloured traces showing predictions from Methods 1–4 respectively. The current measured in Cell #5 in response to Pr6 is shown in grey (same in each row). None of the models were trained on Pr6 data, making this an independent validation. The left panels show the full duration of the signal, and the panels on the right zoom in on selected regions (2.41 to 2.48s, 5.5 to 7.3s, and 7.25 to 7.9s).

A quantitative view of the validation and cross-validation results for Cell #5 is given in Figure 9 (top). The top row of this table shows the RMSE for the independent validation protocol. To enable easier comparisons, the RMSEs have been normalised to the best performing method, so that the best method has relative RMSE 1, while a method with a relative value of 2 has an RMSE that is twice as high. For Cell #5, Method 4 provided the best predictions of the independent test protocol and also performed well on cross-validation. As may be expected, each method had the lowest score on its own fitting data. Data for all 9 cells is given in Supporting Material Figure S13.

**Figure 9:**
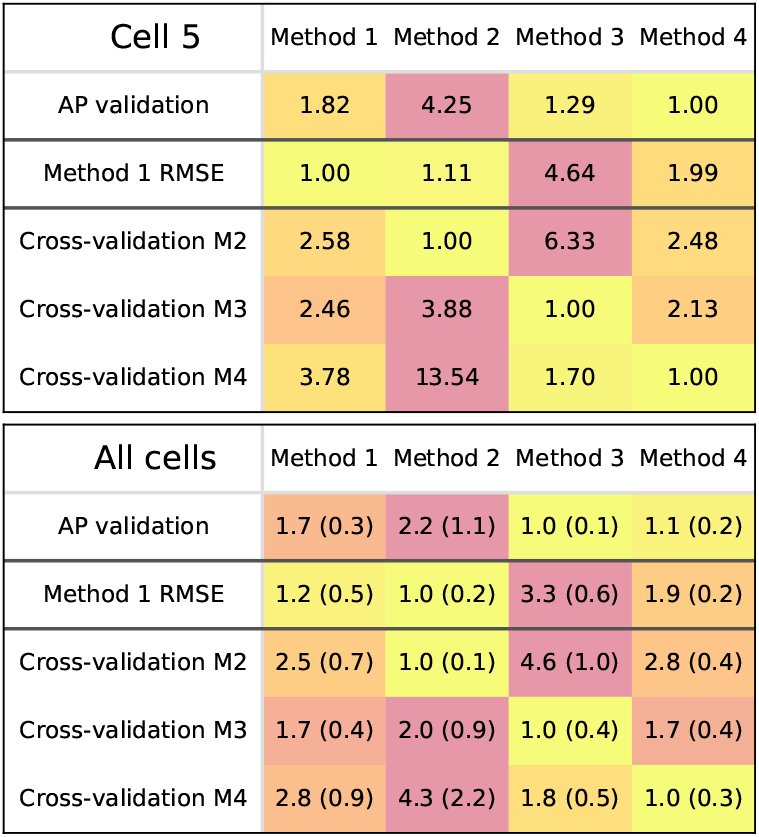
Validation and cross-validation results. (*Top*) Results for Cell #5. Each row shows the relative RMSE for a fitting method, scaled so that the best performing method is indicated by 1, while a method with a relative score of e.g. 1.2 had an RMSE that was 1.2 times larger. The top row shows the relative *E*_AP_ for each method, with the remaining rows showing *E*_M1_, *E*_M2_, *E*_M3_, and *E*_M4_ respectively. (*Bottom*) Mean results over all 9 cells, with standard deviations given in parentheses.

The lower panel in Figure 9 shows similar relative RMSEs, but now presented as the mean and standard deviation for all 9 cells. Here, Method 3 performed best at the AP prediction task, although Method 4 was very close (within standard deviation) and outperformed it at cross-validation.

Interestingly, Method 2 outperformed Method 1 on the *E*_M1_ criterion in the averaged data, and also in 6 out of 9 cells. This indicates the advantage of fitting steady states and time constants simultaneously (as happens in Method 2) over fitting them sequentially (Method 1), as previously described (27, 29). Further illustration is provided in a figure in Supporting Material Section S3.6, where we show that small changes in the slope of the steady state curve have a strong effect on the time constants derived by Method 1; Method 2 can use this to its advantage by slightly adjusting the steady-state curve’s slope to obtain a better fit to time constant data.

### Reliability

Having inspected the predictive capabilities and quality-of-fit of models obtained with each method, we next investigated the reliability of each method’s optimisation procedure. Ideally, a method returns the same result every time it is applied, and indeed Method 1 lives up to this ideal. For Methods 2–4 however, we used (1) randomly sampled initial guesses for parameter sets; and (2) a stochastic optimiser. To increase our chances of finding the best result, we repeated this process 50 times. For a reliable method, we expect a large number of the methods to return similar parameter sets, with similar RMSEs.

Figure 10 gives an indication of each method’s reliability. It shows that Methods 3 and 4 returned consistent RMSEs for at least 20 out of 50 repeats, with the number rising to 40 out of 50 for selected cells. For Method 2, only the first few repeats gave a similar answer, necessitating a larger number of repeats to obtain a reliable result. Method 1 is a deterministic method, and so was omitted from this figure.

**Figure 10:**
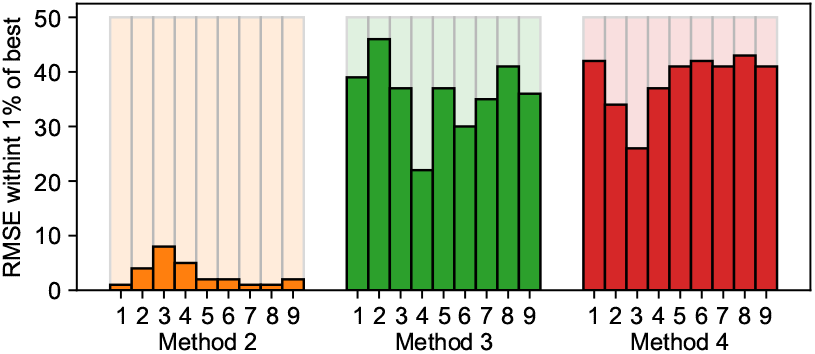
The number of repeats that returned an RMSE within 1% of the best value found in 50 repeats. Data is shown for Methods 2, 3, and 4, for all 9 cells (Method 1 is a deterministic method, and so returns the same answer every time).

To investigate the performance of the different methods, we measured the computation time for each method, and counted the number of forward simulations that were performed. On average, Method 2 was much slower than Method 3, which itself was slower than Method 4 (shown in Supporting Material Figure S14). The number of function evaluations was similar for Methods 3 and 4, indicating that this difference in performance was due to the increased simulation time needed for Method 3 (228s of simulated currents versus 8s of simulation for Method 4). However, many repeats of Method 2 were seen to terminate with a low number of evaluations, which may indicate these optimisations terminated early in a local optimum. To explore this hypothesis, we performed a brute-force exploration of *E*_M2_ and *E*_M4_ for Cell #5, in the region near the optimum returned by Methods 2 and 4 respectively, as shown in Figure 11. For *E*_M4_ we see a clear optimum in each panel, and the function appears smooth (at least within the optimisation boundaries, indicated by the white lines). For *E*_M2_ however, a lot of discontinuities can be seen. The darkest areas in these panels are regions where the intermediate analysis to obtain time constants and steady states failed. As the rightmost panels show, this can occur in otherwise smooth parts of the slices, which may prove challenging for optimisation routines. In addition, a lot of ‘noise’ can be seen (e.g. in the green areas of panels 1), which may indicate the presence of many local minima — although each panel is a 2d slice of a 9-dimensional space, so that this is not necessarily the case.

**Figure 11:**
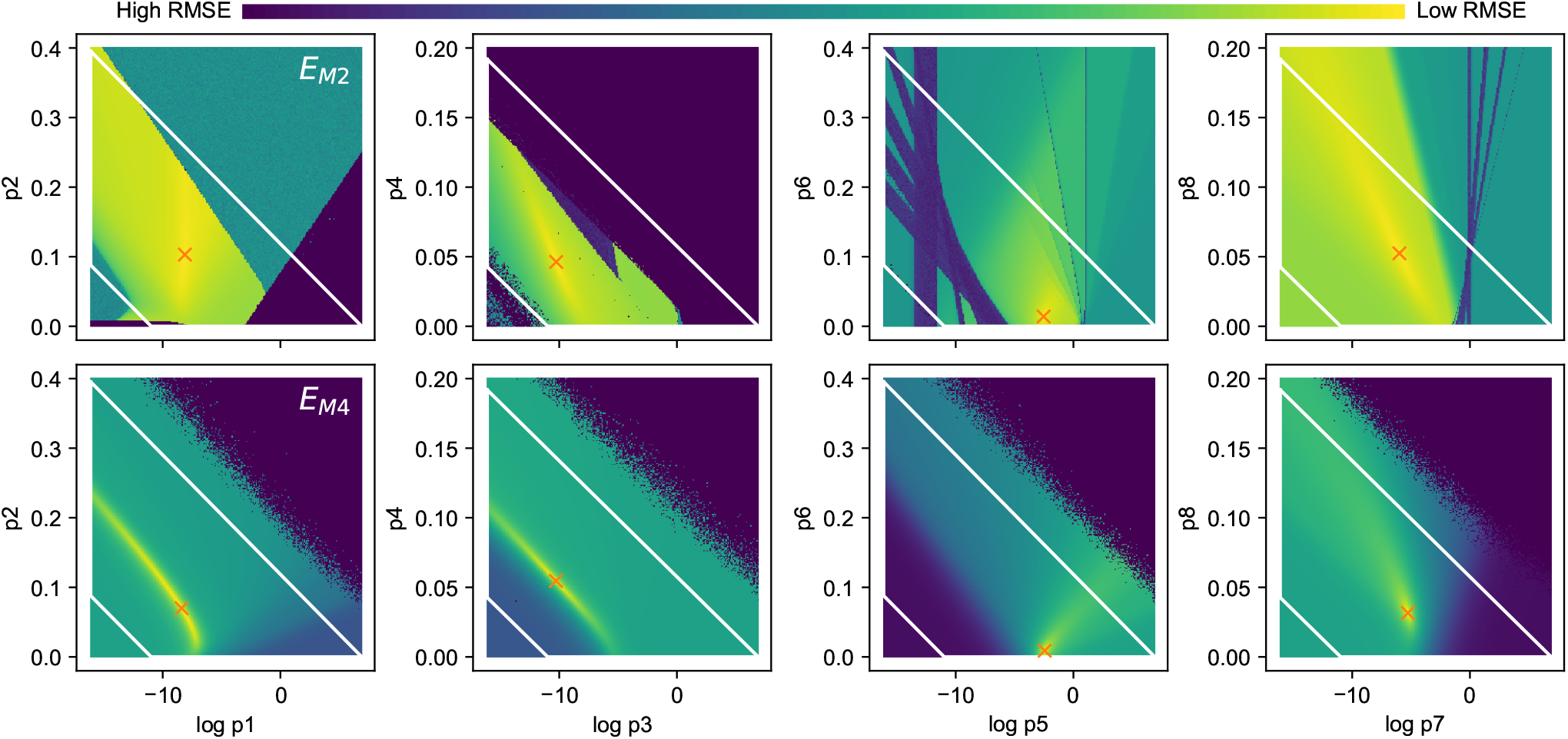
(*Top*) A partial exploration of *E*_M2_ for Cell #5, near the optimum found by Method 2 (indicated by an orange ‘×’). Each panel shows the result of varying 2 parameters, with the other 7 held constant. The parameters varied in the leftmost panels determine the rate of activation, followed by deactivation in the second panel, inactivation in the third and finally the rate of recovery. The white lines indicate the boundaries used during optimisation. (*Lower*) An exploration of *E*_M4_ for Cell #5, near the optimum found by Method 4. The darkest purple colour in the figure represents areas where either summary curve derivation (Top panel) or simulation (Lower panel) failed.

## DISCUSSION

We defined four methods, each representative of a wider class, to fit ion current models using whole-cell current recordings. Methods 3 and 4, both based on whole-current fitting were found to provide the most accurate predictions, while Methods 1 and 2, both based on fitting pre-processed ‘summary’ data, fared more poorly. Of the methods where we applied a stochastic optimisation routine (2, 3, and 4), Methods 3 and 4 were found to provide the most consistent results, and Method 4 was the most time-efficient both in terms of experimental and computational effort.

To further compare the results from different methods, we plotted the fits from each method and each cell in Figure 12. Even in areas where parameter sets overlapped, each method can be seen to introduce its own small bias. The figure also points to a difference in deactivation for Method 3, which consistently placed the deactivation parameters *p*_3_ and *p*_4_ in a different part of the parameter space than the other methods. We found that models using parameter sets found by Method 3 gave the best action potential predictions, although this was very closely followed by Method 4 (see Figure 9). Looking at Figure 6 we can see that many cases where Method 1, 2, and 4 predictions deviated from the measured current were during deactivation. This points to an advantage in describing deactivation for Method 3, and suggests an area where future versions of the sinusoidal protocol used in Method 4 could be improved.

**Figure 12:**
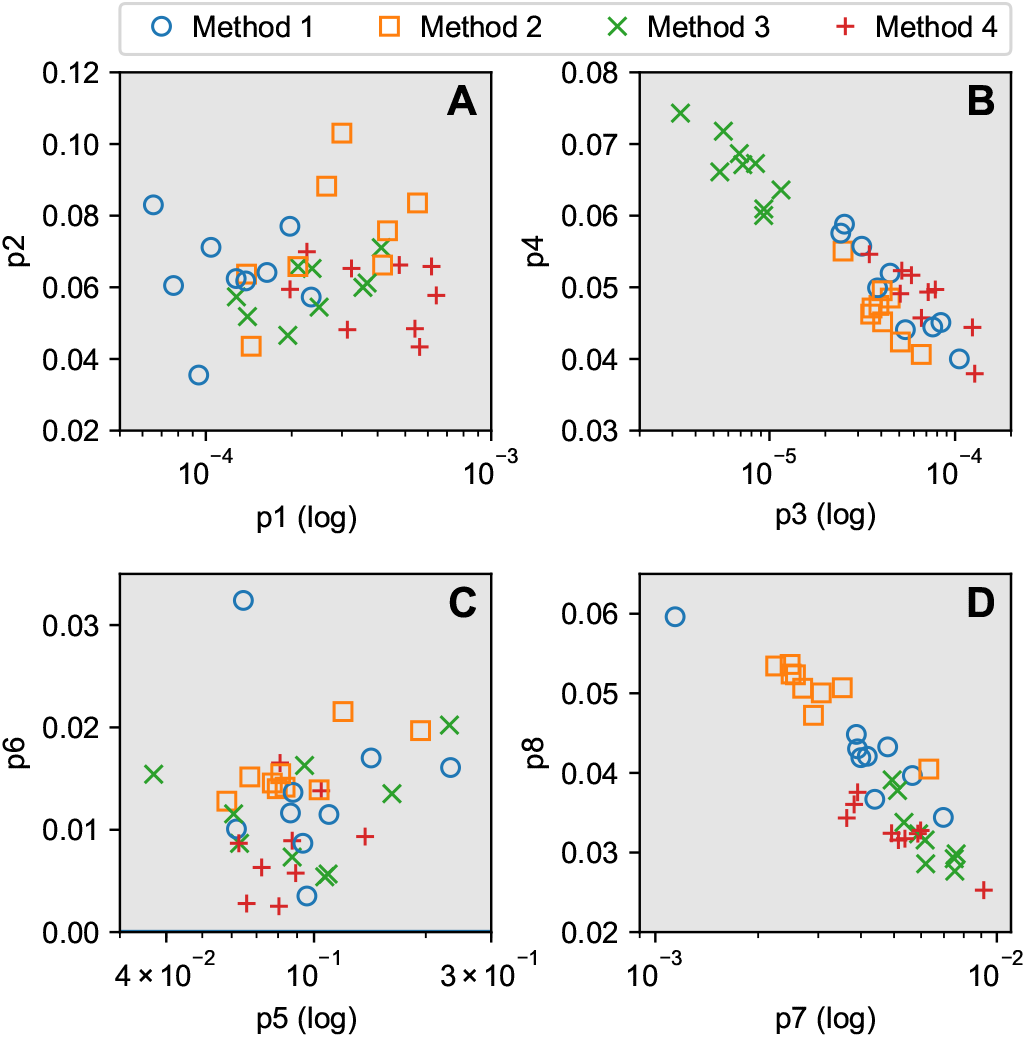
Best solutions returned by the four methods for all cells. The activation rate parameters *p*_1_ and *p*_2_ are shown in panel *A*, deactivation in *B*, and inactivation and recovery in panels *C* and *D* respectively.

For Methods 2, 3 & 4 we observed significant improvements to the performance of optimisation routines by performing a log transform on some of the parameters, providing appropriate constraints on rates and voltage-dependence of the rates, and refining differential equation numerical solver tolerances. At least some of these aspects may address previous findings that whole-trace fitting is difficult (53), as we found very consistent results with an ‘off-the-shelf’ optimiser after adopting these simple approaches.

The optimisation surface for Method 2 is considerably more ragged than Methods 3 and 4, making this a difficult optimisation problem. As Method 2 is very common in the literature on ionic model fitting, this result has some implications for the cardiac modelling community: its reputation as a hard problem requiring special methods may be due to the choice of summary data rather than any intrinsic mathematical or computational difficulty with the models. As the choice of Method 2 is typically based on data availability (43), increased sharing of whole-cell current data (with new information-rich experiments) is a more promising way forward than developing and applying novel optimisation algorithms. For experimenters, as well as universities, journals, and other publishers of scientific data; our findings re-emphasise the need for sharing of raw data, for example in online repositories or data supplements (79).

In cases where full currents are not available (i.e. when using historical data), some steps may be taken to alleviate the problems associated with Method 2. Reviewing Figure 11, some of the high error parts of the *E*_M2_ surface arise from numerical issues during the derivation of summary curves from simulation data (for instance in deriving time constants from almost-flat simulated/experimental current traces). It may be possible to standardise and improve these analysis methods, and so remove some of the discontinuities in *E*_M2_.

Another approach we tried to improve Methods 2 & 3 is to use Method 1 to propose an initial guess parameter set for Methods 2 & 3, instead of sampling a point from randomly within the parameter constraints, as hinted at previously (80). In initial tests, this gave very similar results to the ‘full’ Methods, but with only a single optimisation repeat needed. The full results are shown in Supporting Material Tables S2 & S3.

A reason sometimes given to continue using steady state and time constant curves is that they allow comparison to published data and previous results from the same lab. So Method 3 may be seen to have an advantage over Method 4 in that it contains the traditional protocols needed to derive steady states and time constants. However, as Vandenberg et al. (81) point out, it is important to realise that these values can be highly sensitive to the (occasionally unpublished) details of the protocols and analysis methods, so that making these comparisons is not without danger. We have seen though, that Method 4 can provide excellent simulations of these protocols, so that comparisons to previous data could be made by (1) running an experiment with a novel optimised protocol, (2) quickly and reliably fitting a model to the raw current data, (3) simulating the previously used protocols and performing the analysis. Indeed, this method could be used to compare data sets (via modelling) from protocols with slight discrepancies (82).

### Limitations

In this work we characterised four methods of fitting an ion current model, and proceeded to analyse and critique these methods based on our characterisation. Although we tried not to misrepresent any method, it is therefore important to highlight any areas where our efforts may become subjective or otherwise fall short.

Firstly, running both a full set of conventional protocols, as well as Pr6 and Pr7 in a single cell is experimentally infeasible. For that reason, the study by Beattie et al. (14) used truncated version of the conventional protocols (e.g. with fewer voltage sweeps), and omitted Pr2 variants with different P1 voltages. We must therefore admit the possibility that Methods 1 or 2, performed with a larger dataset, would lead to improved results. In particular, the time constants of activation are ‘missing’ in the range (−30mV, 30mV), which is exactly where the peak value should occur. However, given the excellent performance of Method 3 in our study, it seems the required information was present in the recordings, so that the limited efficiency of both methods still stands.

Another limitation stemming from the cell-specific nature of our measurements is that we cannot benefit from averaging as a noise-reduction strategy. As Figure 2 shows, the mean summary curves over all 9 cells often show qualitative behaviour that is more like the expected results than our individual cell measurements. However, it has also been shown that averaging in cellular electrophysiology can lead to erroneous results (83), and there is a clear benefit of having methods that work on a single-cell, thereby allowing investigations of cell-to-cell variability, heterogeneity, and measurements of cells only available in limited quantities (e.g. human tissue).

In our Method 1, we did not use time-to-peak information (as suggested by (25)), but used what Willms et al. (29) refer to as the ‘disjoint method’ rather than estimating time constants and steady-states simultaneously. We did not, unfortunately, find much mention of such improved methodologies in the applied literature, and so believe that our Method 1 is still a good representation of a commonly followed approach. Similarly, the derivation of the summary curves may be improved in other ways, which would lead to better results for Methods 1 and 2. As a counterpoint, we note that Methods 3 and 4 don’t require these complex intermediate analyses, are easier to implement, and do not rely on the assumption that the summary curves provide an accurate representation of the raw current data.

In deriving Methods 2 and 3, subjective choices had to be made regarding *weighting* of the individual currents, and we accept that changing this may slightly change the results. For example, similar error measures could be derived that do not weight for the length of the protocol, that use fixed scalings per protocol instead of the cell-specific ones we used, or even weight ‘important’ parts of the currents more than others. Method 4 has an advantage here in that there is no such subjective weighting choice to be made.

For Methods 2–4 different error measures could have been used, although most common error measures or log-likelihoods are variants of the sum-of-squares error, and so will share the same optimum (even if the gradients differ, which can affect the optimisation routine). It is tempting to try and improve the methods by carefully crafting new error functions, for example ones that weight rapidly changing areas more strongly than areas with little change, or ones that would emphasise areas known to be important for *I*_Kr_. For the present study, we thought it advisable to avoid such subjective ‘tweaking’, hopefully leading to a result that is representative of standard approaches.

## CONCLUSION

We presented and compared four methods to fit an ion current model, each based on a common class of methods used in the literature. By performing these methods on a set of CHO-cell hERG1a measurements, we found that methods based on whole-current fitting provided both the most accurate and the most reproducible results. Models fit using a novel sinusoidal protocol were found to have similar predictive ability as those fit to conventional voltage-step protocols, while having a lower experimental and computational cost. Further analysis showed that using numerical optimisation to fit a model to steady-state and time-constant data was potentially hazardous, and we point towards increased sharing and use of raw current traces as a viable solution to this problem. Our results highlight some of the remaining challenges to make truly predictive models of ionic currents, and the possibilities for novel experimental protocols to enable faster and more reliable fitting.

## AUTHOR CONTRIBUTIONS

M.C. and G.R.M. designed and performed the analysis. K.A.B. performed the experiments. K.A.B. and G.R.M. designed the experimental protocols. M.C., D.J.G., and G.R.M. wrote the manuscript. All authors approved the final version of the manuscript.

## ACKNOWLEDGMENTS

This work was supported by the Biotechnology and Biological Sciences Research Council [grant number BB/P010008/1]; the Engineering and Physical Sciences Research Council [grant numbers EP/I017909/1 and EP/K503769/1]; and the Wellcome Trust [grant numbers 101222/Z/13/Z and 212203/Z/18/Z]. M.C., G.R.M. and D.J.G. acknowledge support from a BBSRC project grant. K.A.B. was supported by the EPSRC via PhD studentship and postdoctoral support. G.R.M. acknowledges support from the Wellcome Trust & Royal Society via a Sir Henry Dale Fellowship and a Wellcome Trust Senior Research Fellowship. K.A.B. is an employee and shareholder of GlaxoSmithKline Plc.

## Footnotes

1 Note that the independence assumption means that, somewhat confusingly (but perhaps appropriately), channels can be both ‘activated’ and ‘inactivated’ at the same time, as the opposite of activation is called deactivation and the opposite of inactivation is recovery.

