## Supporting Material for "Four ways to fit an ion channel model"

This document contains supporting material for the article “Four ways to fit an ion current model”. Further figures, animations, and code can be found at `https://github.com/CardiacModelling/FourWaysOfFitting`.

### S1 UNDERSTANDING VOLTAGE PROTOCOLS

In this section we analyse the protocols used in this study (and in Beattie et al. (1)), using phase plane analysis. The first four protocols, Pr2–5, are adaptations of common voltage clamp protocols used to characterise $I_{\mathrm{Kr}}$, while Pr7 is a novel sinusoidal protocol intended to provide the same information in a much shorter time. Pr6 is a collection of (regular and irregular) action potential wave forms to measure the behaviour of $I_{\mathrm{Kr}}$ under physiological and pathological conditions.[1] As in Beattie et al. (1), we used Pr6 as a *validation* protocol, while either Pr7 or the set Pr2–5 were used for model *fitting*. Note that the full set of protocols was run on every cell.

There are some similarities between the protocols. Pr2–5 are all periodic protocols, repeated several times (with each repeat shown in a different colour in the figures) with a change either in one step's duration (Pr2) or voltage (Pr3–5). All protocols in this study start with a constant holding potential of −80 mV followed by a brief step down −120 mV. Because $I_{\mathrm{Kr}}$ is mostly inactive at these potentials, this allows the $I_{\mathrm{Kr}}$-independent leak current to be estimated and subtracted from the signal (1). The protocols end with another step down to −120 mV, which is intended to rapidly bring the channels into a closed state, thereby reducing the time needed to settle back to steady state between repeats or between experiments.

#### S1.1 Phase plane analysis

Using a two-dimensional model (see main manuscript) allows us to represent each possible state as a point on a *phase plane*, in which we plot activation $a$ on the x-axis and recovery $r$ on the y-axis. By running simulations and plotting the trajectories of $a$ and $r$ in the plane we can show the motivation behind different voltage-clamp protocols in terms of the types of behaviour they provoke.

Held at any given voltage $V$, the model will eventually converge to a steady state $a = a_\infty(V)$, $r = r_\infty(V)$ known as a *stable node*, see Figure S1.A. When $V$ is changed abruptly, the stable node instantaneously moves to a new position, to which the states then converge with speeds dictated by $\tau_a(V)$ and $\tau_r(V)$. With typical parameters for $I_{\mathrm{Kr}}$, inactivation/recovery is orders of magnitude faster than activation/deactivation so that many trajectories through the phase space will start with fast vertical movement followed by a slower horizontal drift. The model conductance at any point $(a, r)$ in the plane is proportional to the product $a \cdot r$. Connecting points with an equal $a \cdot r$ leads to the iso-conductance lines shown in Figure S1.B, which indicate the fraction of maximal conductance $g_{\mathrm{Kr}}$ in different regions. Using these two graphs an intuitive idea of $I_{\mathrm{Kr}}$ behaviour may be developed, and we will use these phase planes to analyse different voltage protocols throughout the text. An example of a phase plane trajectory is shown in Figure S2.

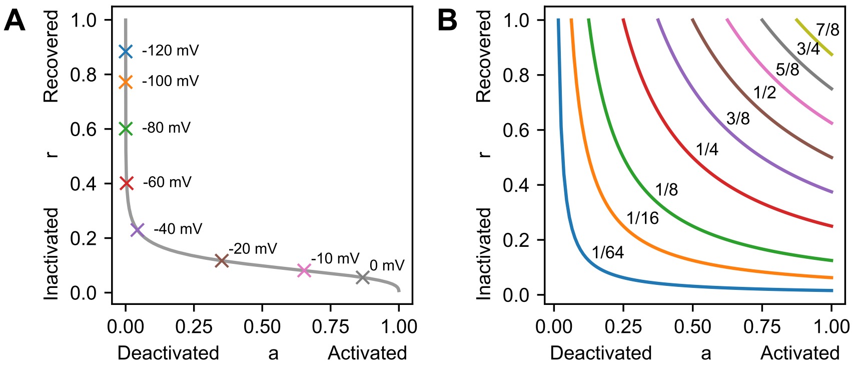

Figure S1: A guide to interpreting phase portraits. (*A*) At any voltage $V$, the point $(a_\infty(V), r_\infty(V))$ forms a *stable node* to which the system, if held at this $V$, will converge. The grey line in the figure is formed by plotting these stable nodes for a wide range of voltages, based on a simulation with the parameters for Cell #5 identified in Beattie et al. (1). (*B*) Different points in the phase plane correspond to different fractions of the maximal conductance (given by the product $a \cdot r$). When all the points for a given fraction are connected they form the iso-conductance lines shown here. Combining these two figures we can see that the channel's steady-states have low conductance while any larger currents are necessarily transient.

[1] Although we measured current through a hERG1a channel in a CHO cells, we will occasionally use the shorthand $I_{\mathrm{Kr}}$ to describe the current in this manuscript

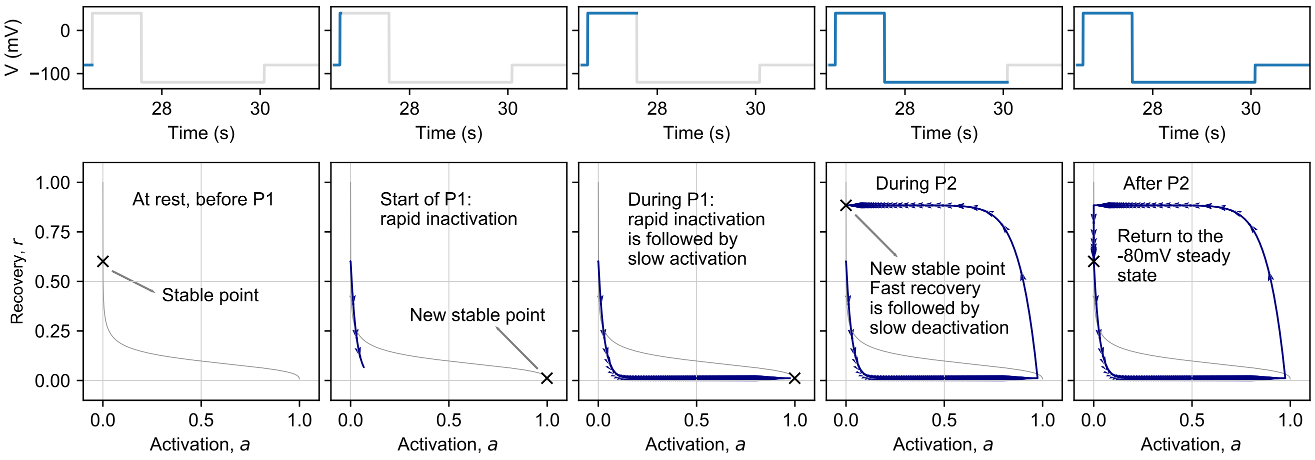

Figure S2: A simulated phase plane trajectory for the final repeat of protocol Pr2. In the first panel, the cell is being held at −80mV, and the system is in its steady state $a_\infty(V = -80), r_\infty(V = -80)$. In the next panel, a voltage step (P1) is applied, causing an instantaneous jump of the stable point towards the bottom right of the graph. The system now starts to rapidly inactivate, resulting in a downward trajectory in the phase plane. After a few milliseconds, inactivation is nearly complete, and activation begins to dominate the trajectory, resulting in the horizontal trajectory shown in the third panel. At the end of P1, the system is at (or very close to) its new stable state. Now, a new step (P2) is applied, again leading to an instantaneous jump of the stable state, followed by a vertical-then-horizontal trajectory of the system through phase space. In the final panel, the original membrane potential is restored, causing the system to revert to its original stable state.

#### S1.2 Pr2: A time constant of activation

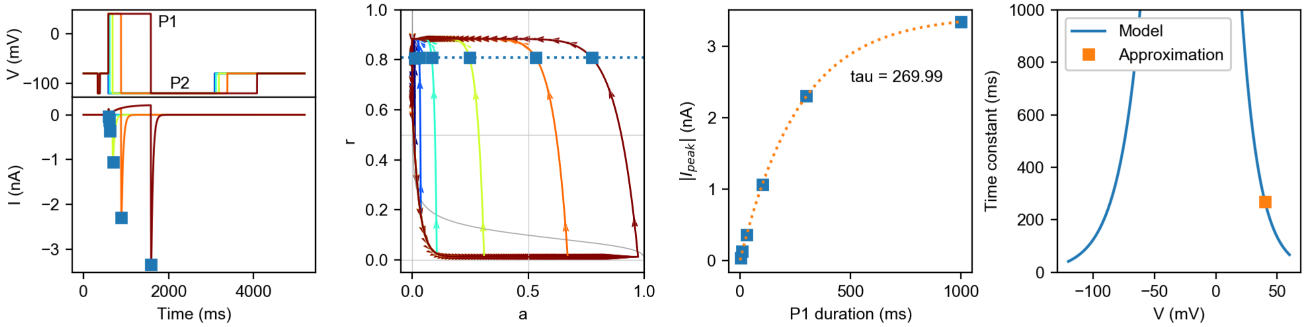

Figure S3: Simulated analysis of Pr2, approximating the time constant of activation at $V_2 = 40$mV. The protocol and current are shown in the left-most panel, with the peak currents during each repeat highlighted. The same highlighting is applied in the phase diagram, which shows that all peaks occur at almost the same level of recovery, $r \approx \tilde{r}$. Next, the peak current is plotted against the P1 duration and fit with a single exponential. This results in a single time constant, which is shown in the final panel along with the underlying model variable.

Pr2 (6 repeats of 5.2s each, 31.2s in total) is used to obtain an approximation of the time constant of activation at $V = 40$ mV. Its main feature is a variable-duration step (P1) at +40 mV. In the phase diagram this corresponds to a movement from top-left (the steady-state for −80 mV) down to the stable node for +40 mV in the lower-right part of the plane. Note that only the longest step (darkest red) actually reaches the stable node, while in the other repeats the step ends before the it is reached. During this time, the model is inactivated ($r \approx 0$) to approximately the same degree for each repeat, while the activation level varies depending on the time spent at 40 mV — it is this activation level that we wish to measure. To that end, P1 is followed by a step (P2) down to −120 mV, triggering a rapid recovery and a measurable current.

We now inspect the peak currents during P2, $I_{\text{peak}}$. From the phase diagram we can observe that the level of recovery at the peak is roughly the same for each repeat, so that we can approximate it by some constant (but unknown) value $\tilde{r}$, and write

$$I_{\text{peak}} \approx g_{\text{Kr}} \cdot a(t_{\text{peak}}) \cdot \tilde{r} \cdot (V_2 - E_K), \tag{S1}$$

where $V_2$ is the voltage during the P2 step, and $a(t_{\text{peak}})$ is some unknown activation level. We can also see that the trajectory from the end of P1 (a point near the x-axis) up to the point where peak current occurs (blue squares) is near-vertical, so that the level of activation at the peak, $a(t_{\text{peak}})$, is approximately equal to the level at the end of P1, $a_1$, so that we can write

$$I_{\text{peak}} \approx g_{\text{Kr}} \cdot a_1 \cdot \tilde{r} \cdot (V_2 - E_K), \tag{S2}$$

Next, we solve the differential equation for $a$ under a fixed voltage, to find

$$a(t) = a_\infty(V) - (a_\infty(V) - a_0)e^{-t/\tau_a(V)} \tag{S3}$$

where $V$ is the voltage during the step, $t$ is the time since the start of the step, and $a_0$ is the level of activation at the start of the step. Adapting this for the activation at the end of P1, we fill in $V = V_1$ and set $t$ equal to the step duration $t_1$ to find

$$a_1 = a_\infty(V_1) - (a_\infty(V_1) - a_0)e^{-t_1/\tau_a(V_1)} \tag{S4}$$

which we combine with the equation for $I_{\text{peak}}$ to find

$$I_{\text{peak}} \approx g_{\text{Kr}} \cdot \tilde{r} \cdot (V_2 - E_K) \cdot \left(a_\infty(V_1) - (a_\infty(V_1) - a_0)e^{-t_1/\tau_a(V_1)}\right) \tag{S5}$$

$$= c_1 + c_2 e^{-t_1/\tau_a(V_1)}. \tag{S6}$$

In other words, $I_{\text{peak}}$ should (approximately) be a function of the P1 duration $t_1$ and three unknowns $c_1$, $c_2$, and $\tau_a(V_1)$. As a result, we can plot $I_{\text{peak}}$ against the P1 duration $t_1$, and fit a single exponential to find the time constant $\tau_a(V = V_1)$.

In a typical run of experiments, this protocol would be repeated with different P1 voltages, resulting in time constants for several voltages. As the data set from (1) does not include these, we will instead obtain further time constants of activation from Pr5. Finally, a slight variation of this protocol was used in cells 7 and 8, details of which can be found in the code published with this manuscript at https://github.com/CardiacModelling/FourWaysOfFitting.

#### S1.3 Pr3: The steady state of activation

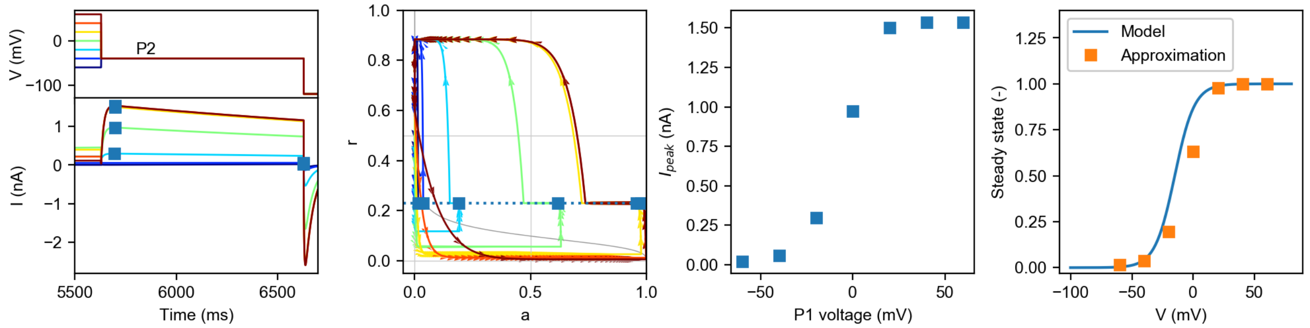

Figure S4: Simulated analysis of Pr3, approximating the steady state activation curve. The protocol and current are shown in the left-most panel, with the peak currents during each repeat highlighted. The same highlighting is applied in the phase diagram, which shows that all peaks occur at almost the same level of recovery. Next, the peak current is plotted against the P1 voltage. Because each repeat had approximately the same recovery level, $V = V_2$ for each repeat, and because the highest voltage leads to $a_\infty \approx 1$, we can normalise the peak currents by dividing through the highest obtain value to find the approximation of the steady state of activation shown in the final panel. Note the difference between the true model variable and the approximation, which is due to the incomplete approach to the steady state for voltages around 0 mV, which can be seen in the phase diagram.

Pr3 (7 repeats of approx. 8.3s each, 58s in total) is intended to characterise $I_{\text{Kr}}$'s steady state of activation for several voltages. Its main feature is a 5 s long variable-voltage step, P1, followed by a step P2 down to a fixed voltage, during which current is measured. In the phase plane, P1 is visible as a downwards movement from the steady state at −80 mV (top left for all repeats) to a new steady state lower on the plot. This steady state is close to $a = 1$ for high positive voltages (but notice that lower voltages have not yet quite reached the stable point after 5 s, which will be important later). Due to the large difference in the time constants of activation and recovery the P1 currents show a rapid downward movement, followed by a slower horizontal drift. At the end of P1, the system is close to the steady state of activation for the tested voltage, so that measuring the current at this point in time would provide us with clear information about the voltage-dependence of activation. Unfortunately, the level of inactivation at this point makes this current very small, so that it cannot be measured with a reasonable signal-to-noise ratio. The P2 step to a fixed voltage of −40 mV elicits a much stronger $I_{\text{Kr}}$ current by causing rapid recovery from inactivation. This is visible in the phase plane as a rapid upward movement, which abruptly stops and turns into a slow leftward drift. Two interesting things happen near this abrupt 'corner' in the graph: (1) as this point is the furthest top-right of any point in P2, this is where the P2 peak current occurs; (2) recovery reaches its steady-state for the P2 voltage ($r_\infty(V_2) \approx 0.23$), which is the same for every test voltage repeat. Since there has been very little change in activation $a$ when the peak current is reached, we can approximate the peak current by

$$I_{\text{peak}}(V_1) \approx g_{Kr} \cdot a_\infty(V_1) \cdot r_\infty(V_2) \cdot (V_2 - E_K) \tag{S7}$$

where $V_1$ and $V_2$ are the voltages during P1 and P2 respectively. At the highest voltage tested $V_1 = V_{\text{max}}$, we can assume that $a_\infty(V_{\text{max}}) \approx 1$, so that we can write

$$I_{\text{max}} = I_{\text{peak}}(V_1) \approx g_{Kr} \cdot r_\infty(V_2) \cdot (V_2 - E_K) \tag{S8}$$

As a result, we can divide by $I_{\text{max}}$ to obtain

$$\frac{I_{\text{peak}}}{I_{\text{max}}} \approx a_\infty(V_1). \tag{S9}$$

This can be plotted to give the summary curve shown in Figure S4, and is commonly known as the 'activation curve'.

#### S1.4 Pr4: Time constant of inactivation

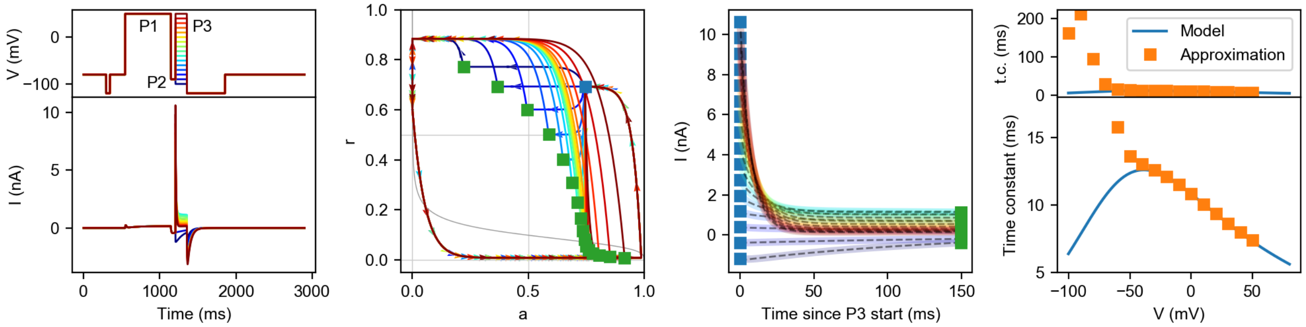

Figure S5: Simulated analysis of Pr4, approximating the time constant of inactivation. The protocol and current are shown in the left-most panel. Next, a phase diagram is shown with the start of P3 highlighted with a blue square, and the end of P3 for every repeat shown with a green square. Simulated currents during P3 are shown in the third panel, and are well-fitted by a single exponential. This yields the time constants shown in the final panel. The values obtained this way are accurate for higher voltages but contaminated by activation for lower voltages. Looking back to the phase diagram, the low voltages (blue lines) have trajectories with a strong horizontal (activation) component, while only the higher voltages (dark red lines) have trajectories determined mostly by recovery.

Pr4 (16 repeats of approx. 2.9s each, 46s in total, can be used to approximate the time constant of inactivation, and consists of a long step (P1) at +50 mV, followed by a quick step (P2) down to −90 mV and finally a variable-voltage step P3. During P1 the model quickly inactivates and then activates, which is visible in the phase plane as a movement from top-left down to the lower-right corner. P2 then causes a rapid recovery (and a large current), and only minor deactivation: an upwards movement in the phase plane that is deflected left, to a point near the phase-plane coordinates ($a = 0.75$, $r = 0.7$).

Next, the short variable-voltage step P3 is applied. Since P2 left the channels both activated and recovered, large currents can be recorded throughout P3. Due to P3's short duration and the large difference between the rates of activation and inactivation, the decays of most of these currents are characterised almost entirely by inactivation (vertical versus horizontal movements on the phase plane). As a result, we can fit exponential curves to these decays to approximate a time constant of inactivation for every tested voltage. Note however, that this assumption is increasingly invalidated for lower potentials, so that Pr4 can only be used to approximate time constants for higher voltages. The resulting time constant approximations are shown in Figure S5.

Note: Some of the most striking parts of the phase plane diagram for Pr4 correspond to the step *after* the very short P3. To trace the movements of the system through the phase plane it may be helpful to consult the 3-dimensional phase diagrams given in Figure S10 or the videos at `https://github.com/CardiacModelling/FourWaysOfFitting`.

#### S1.5 Pr5: Time constants, IV curve, and steady state of inactivation

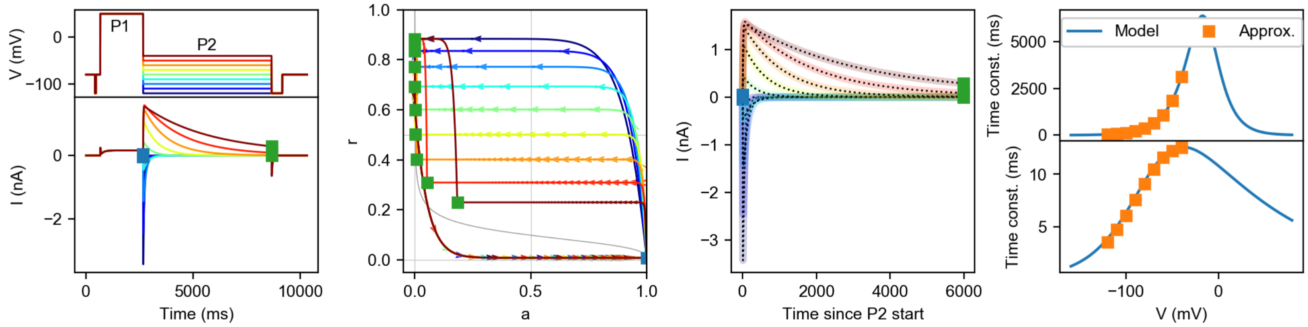

Figure S6: Simulated analysis of Pr5, approximating the time constants of activation and inactivation. The protocol and current are shown in the left-most panel, with the start and end of the P2 step indicated with blue and green squares respectively. The same highlighting is applied in the phase diagram, which shows all P2 currents start with rapid recovery, followed by slow deactivation. In the next panel, the P2 currents are plotted and shown to be well fitted by a double exponential. This results in two time constants, which are shown in the final panels along with the underlying model variables.

Pr5 (9 repeats of approx. 10.3s each, 93s in total) is used to estimate time constants of both activation and inactivation, as well as providing a graph of voltage-dependent peak currents (the ‘IV curve’) from which the steady states of inactivation can be approximated. The main part of Pr5 consists of a step to +50 mV (P1) followed by a variable-voltage step (P2). As before, the P1 step of Pr5 can be seen in the phase plane as a movement from top-left to a stable point in the lower-right of the plane where the channels are almost entirely activated ($a \approx 1$) and inactivated ($r \approx 0$). Next, a much lower voltage is applied during P2, causing the channels to rapidly recover and resulting in a strong current. This is shown as an upward movement in the phase plane, which then gradually turns into a horizontal movement as the system begins to deactivate. As a result, P2 is characterised by a very rapid deflection (positive or negative depending on the sign of $V_2 - E_K$) caused by recovery-from-inactivation, followed by a much more gradual decay as deactivation sets in.

We can use this two-phase character of the current by fitting one exponential to the start of the P2 current (pre-peak) to estimate the time constant of inactivation ($\tau_r$), and fitting a second exponential to the end of the current (post-peak) to estimate the time constant of activation ($\tau_a$) for each test voltage $V_2$. These estimates can be improved by fitting the sum of both exponentials directly to the currents, using the independently acquired values as initial guesses. The resulting values are plotted in Figure S6.

#### IV Curve and steady state of inactivation

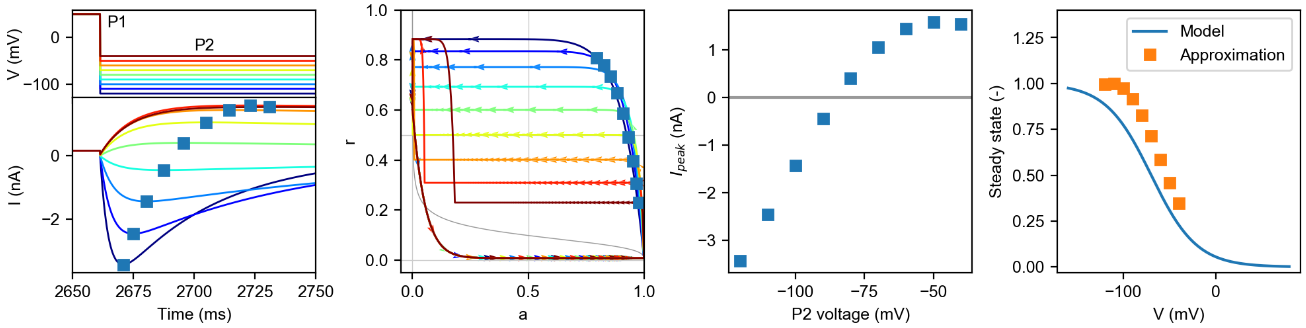

Figure S7: Simulated analysis of Pr5, approximating the steady state of inactivation. The protocol and current are shown in the left-most panel, with the peak currents during each repeat highlighted. The same highlighting is applied in the phase diagram. Next, the peak current is plotted against the P2 voltage, resulting in a (commonly referred to as 'the') IV curve. By dividing the peak current through $V_2 - E_K$ we obtain $g_{\mathrm{Kr}} \cdot a(t) \cdot r(t)$. We then use the approximations $a(t) \approx 1$ and $r(t) \approx r_\infty(V_2)$, to find $g_{\mathrm{Kr}} \cdot a(t) \cdot r(t) \approx g_{\mathrm{Kr}} \cdot r_\infty(V_2)$. Finally, we assume that the peak $r$ measured is $\approx 1$, so that $g_{\mathrm{Kr}} \approx \max\left[g_{\mathrm{Kr}} \cdot r_\infty(V_2)\right]$, and we can divide by this value to find $r_\infty(V_2)$ for every tested $V_2$. The resulting values are shown in the final panel, and can be seen to differ from the underlying model variable. Looking at the phase diagram, we see that the final assumption (peak $r \approx 1$) does not hold, but also that low-voltage peaks do not occur exactly at $r(t) = r_\infty$.

In the second application of Pr5, the steady-state of inactivation ($r_\infty$) is approximated in a similar manner to Pr3. First, we extract the peak current during P2 and plot it as a function of voltage. The result is known as an IV curve, and is shown in Figure S7. Again, note that the peak current occurs when the trajectory in the phase plane changes from vertical (upwards) to horizontal (leftwards) movement, and that — especially for the higher voltages — there is relatively little deactivation at this point. As a result, the peak current can be approximated as

$$I_{\mathrm{peak}} \approx g_{\mathrm{Kr}} \cdot a_\infty(V_1) \cdot r_\infty(V_2) \cdot (V_2 - E_K) \qquad \text{(S10)}$$

where $(V_2 - E_K)$ is known and $a_\infty(V_1) \approx 1$ at +50 mV. Dividing by these two quantities, we find an approximation for $g_{\mathrm{Kr}} \cdot r_\infty(V_2)$. If we further assume that for the lowest voltage $r_\infty(V_{\mathrm{min}}) \approx 1$ (again, note for later that it is actually closer to 0.9 in this parameterisation of the model) we can divide by $g_{\mathrm{Kr}} \cdot r_\infty(V_{\mathrm{min}})$ to find an approximation of $r_\infty(V_2)$ for every tested $V_2$. The result is shown in Figure S7

##### S1.5.1 A note on calculating steady-state of inactivation

Calculating steady states of activation and inactivation requires a division by $(V - E_K)$, creating a singular point at $V = E_K$ where conductance cannot be calculated, but more importantly a region around the point $V = E_K$ where any small error is amplified. Looking at Figure S7, we can see that many of the most rapidly changing (and therefore most informative) parts of the inactivation curve occur in this region.

In Figure S8.A, we have plotted the peak currents during the P2 step of Pr5 for all cells. A clear and regular trend can be seen for each cells, and all cells show qualitatively similar behaviour. In the next panel (Figure S8.B) we show the multiplication factor $(V - E_K)^{-1}$ (blue line) that is applied to the panel A data to obtain the steady-state curve (using $E_K = -88.4$mV). To illustrate what will happen if the term $V - E_K$ is imperfectly known we also plot the 10-th and 90-th percentile of the distribution $1/\mathcal{N}(V - E_K, \sigma)$, where $\mathcal{N}$ is a normal distribution and a (somewhat arbitrary) estimate $\sigma = 2$mV. This type of error could easily arise if the calculated $E_K$ differs from the true reversal potential, or if the true transmembrane voltage differs from the command potential.

The summary curves for all cells in Figure S8.C show that this is not just a hypothetical concern, with most cells showing a dramatic deviation at -90mV (and even a change of sign for cells 5 and 6). As a result, we had to omit the data from $V = -90$mV from the summary curves for the steady state of inactivation. In Figure S8.D we plot the same data with a rescaled y-axis, and omit the -90mV points, but now it becomes clear that the wide region of error predicted by panel B is also borne out in practice, as several of the point at -100mV and -80mV also show a strong deviation from the expected sigmoid voltage-dependence.

In the summary curves used in this work, we omitted the -90mV point for both the steady state of inactivation and the time constant of inactivation (where nearness to -90mV caused problems when fitting an exponential curve). Finally, we point out that these issues (caused in part by experimental noise) can be somewhat reduced by averaging the values for all cells (a common noise-reduction technique): the mean value shown is not perfect, but displays a clearer sigmoid form than the data for the individual cells. This means that this issue, although present in all studies that use a similar summary curve, becomes more apparent when aiming for cell-specific results (see also (2)).[2]

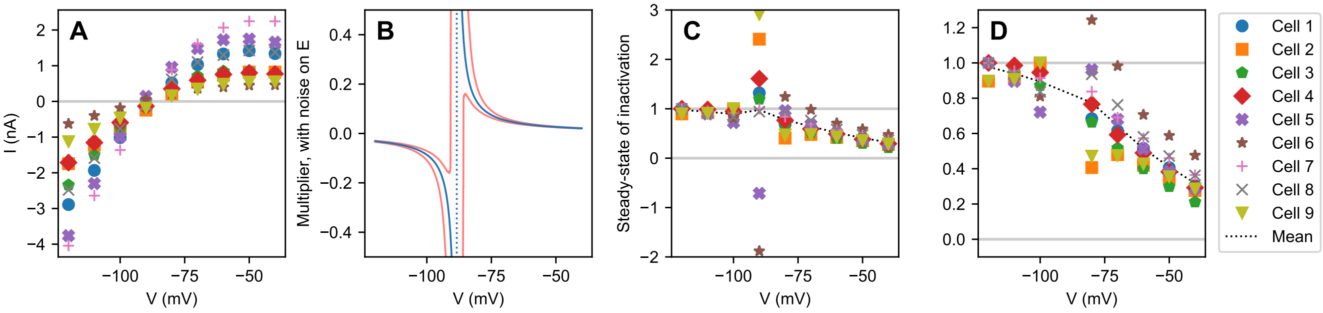

Figure S8: Approximating the steady state of inactivation. (*A*) The calculation of the steady state of inactivation starts from the IV curve data from Pr5. This data is smooth, and has the same qualitative nature for every cell. (*B*) Next, the data is multiplied by a term $1/(V - E_K)$. As shown by the blue line, this term has a large magnitude near $V = E_K$, resulting in a strong amplification of measurement error. The effect of small errors can also be seen by the red lines, which indicate the 10-th and 90-th percentile of the distribution $1/\mathcal{N}(V - E_K, 2\text{mV})$. (*C*) As expected, a major disruption is visible in the experimental data, especially near $V = -90\text{mV} \approx E_K$. (*D*) Omitting the data points for $V = -90$mV removes the largest errors, but strong effects can still be seen for $-100$, $-80$, and $-70$mV.

[2] It may be possible to deal with this issue by introducing a suitable noise model, and performing a weighted fit that assigns lower importance to the affected points. However, as we did not see this approach in our literature review we did not include such a method in this work.

#### S1.6 The summary curves don't match the model variables

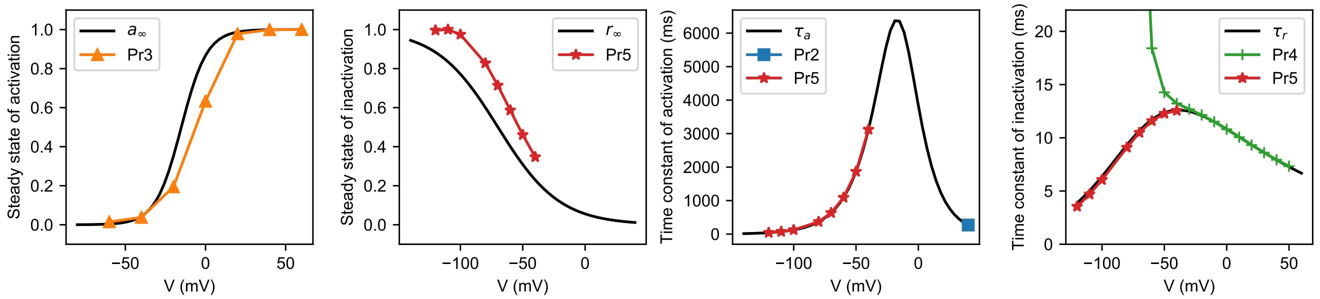

Figure S9: (*Black lines*) The model variables $a_\infty$, $r_\infty$, $\tau_a$, and $\tau_r$, all drawn using the parameters for Cell #5 given in Beattie et al. (1). (*Coloured lines*) Simulated summary curves, obtained by using the same model parameters, running simulations for Pr2–5 and performing exactly the same analysis as on the experimental data. The protocol from which each data point originates is indicated in the legend.

In explaining the rationale behind Pr2–5, we have relied heavily on the idea that they are designed to approximate the model variables $a_\infty$, $r_\infty$, $\tau_a$, and $\tau_r$. Historically, this certainly seems to how these protocols originated (see e.g. Hodgkin and Huxley (3)). However, as theories and models of $I_{\text{Kr}}$ (and other currents) have grown in complexity the strong connection of protocols like Pr2–5 to their modelling origins has increasingly been lost. The analysis methods have, however, remained important as *biomarkers* in their own right, and many physiologists (aware of the shortcomings and pitfalls we discuss below) have chosen to interpret them as such. However, even if we take this view, the best *interpretation* we have of what these biomarkers signify is still that they resemble the variables of a two-state Hodgkin-Huxley model fit to the data. In addition, Method 1 relies on the assumption that these procedures accurately approximate the model variables, so that it's worthwhile pointing out some of the issues in their calculation below.

**Steady-state of activation:** The central idea of Pr3 was to reach the steady state of activation (i.e. reach the stable point for the P1 voltage) and quickly measure a current during P2. Inspecting the phase diagram for Pr3 we can see that this objective is not met for the voltages around 0mV (e.g. the blue and green lines). This leads to an underestimation of the activation at these voltages, which causes the rightward shift of the estimated steady state of activation observable in Figure S9. Less clear from this graph, is that the voltage-dependency of the effect will have caused a change in the *slope* of the estimated steady state curve. Note that using a longer variable voltage step would have brought the system closer to the stable point and reduced the apparent shift, while a shorter step would have caused an even stronger rightward shift. This time-dependency has been recognised by e.g. Vandenberg et al. (4), who warn only to compare data from protocols with an equal P1 duration, which they term 'isochronal activation data'. Similar issues have been recognised for other currents. For example, in a 1992 publication on $I_{\text{Na}}$, Sakakibara et al. (5) consistently avoid the term 'steady state of activation' in favour of 'normalized conductance-voltage relation', which much more cautiously describes what the 'activation' protocol has actually measured.

**Steady-state of inactivation:** The method to obtain a steady-state of inactivation from Pr5 relied on the assumptions that (1) the peak currents measured were not strongly affected by deactivation and (2) that the lowest voltage tested induced an inactivation level $r_\infty(V_{\text{min}}) \approx 1$. As can readily be seen from the phase diagram, the first assumption is violated increasingly at lower P2 potentials. This leads to both an underestimation of the steady state of inactivation (visible as a rightward shift) and a change in the apparent slope of the inactivation curve (as lower voltages are affected more than higher ones). Looking back at the phase diagram, it is clear that the second assumption is violated too, and as a result the normalisation of the estimated curve will be off, again leading to changes in both midpoint and slope of inactivation. This second problem could be perhaps addressed by adding even lower potentials to the protocol, but notice that this would exacerbate problem 1. Staying with Pr5, it seems from the phase diagram that for lower voltages the separation between the recovering and deactivating parts of the P2 current becomes increasingly less clear. However, the strategy of fitting both at once appears to have paid off, as the Pr5 time constants in both right-hand panels of Figure S9 overlap well with the model variables.

**Time constants:** Finally, as already noted Pr4 fails to estimate good time constants for lower potentials (see the right-most panel in Figure S9, but this limitation can be overcome by using the Pr5 derived points instead. However, the curves from both protocols don't quite line up, indicating further issues with one or both analyses.

**Previous work:** Please note that the above demonstrations are not novel, but reaffirmations of (much) earlier work by e.g. Beaumont et al. (6, 1993), Willms et al. (7, 1999), and Lee et al. (8, 2006).

#### S1.7 Pr6: AP validation protocol

An action potential voltage clamp protocol (Pr6) is used in this study and in Beattie et al. (1) as a *validation protocol*: instead of fitting to data from these measurements we use it to evaluate predictions from models fit to the other data sets. Unlike Protocols 2–5, Pr6 does not contain any repeats. The bulk of Pr6 is a sequence of realistic action potentials, so that validation happens against physiologically relevant situations including pathological after-depolarisations. Looking at the phase diagram in the main paper (which has been coloured through time to match Pr6's voltage and current traces) we can see a secondary effect: the short time between the APs causes a build-up of activation, so that the early parts of Pr6 are a proxy for $I_{\mathrm{Kr}}$ during low heart rates, while the latter parts elicit $I_{\mathrm{Kr}}$ behaviour during periods of prolonged higher rates.

#### S1.8 Pr7: Sinusoidal protocol

Pr7 (single run of 8s) is a novel sinusoidal protocol introduced in Beattie et al. (1). Like Pr6, it does not contain any repeats but instead consists of a single eight second sweep. The protocol starts with a step to +40 mV followed by a step down to −120 mV, eliciting high conductance and a strong negative current. In the phase diagram shown in the main paper, this corresponds to the blue trajectory from lower-right to top-left. Note how the blue line goes close to the top-right corner of the plane (high $a$ and $r$). This appears to be crucial in the protocol design for estimating the conductance parameter $p_9 = g_{\mathrm{Kr}}$ accurately: at the point $(1, 1)$ in the phase plane the current would be given by $I_{\mathrm{Kr}} = g_{\mathrm{Kr}}(V - E_K)$ allowing $g_{\mathrm{Kr}}$ to be estimated directly. By including a step that approaches this point, we gain a lot of information about $g_{\mathrm{Kr}}$.

The remainder of the protocol consists of three sine waves of varying frequencies added together. In the phase diagram, this induces rapid near-vertical movements from which we can infer the properties of inactivation, but also slower horizontal movements that tell us about activation, across the full physiological range of voltages. Importantly, many changes in the trajectory happen far from the x and y-axis – in other words – many of the dynamical changes induced by Pr7 occur while strong $I_{\mathrm{Kr}}$ is being generated and the current is experimentally observable.

#### S1.9 Three-dimensional phase diagrams

The voltage-dependence of the steady state can be made more clear by plotting the phase diagrams in three dimensions, with voltage on the third axis.

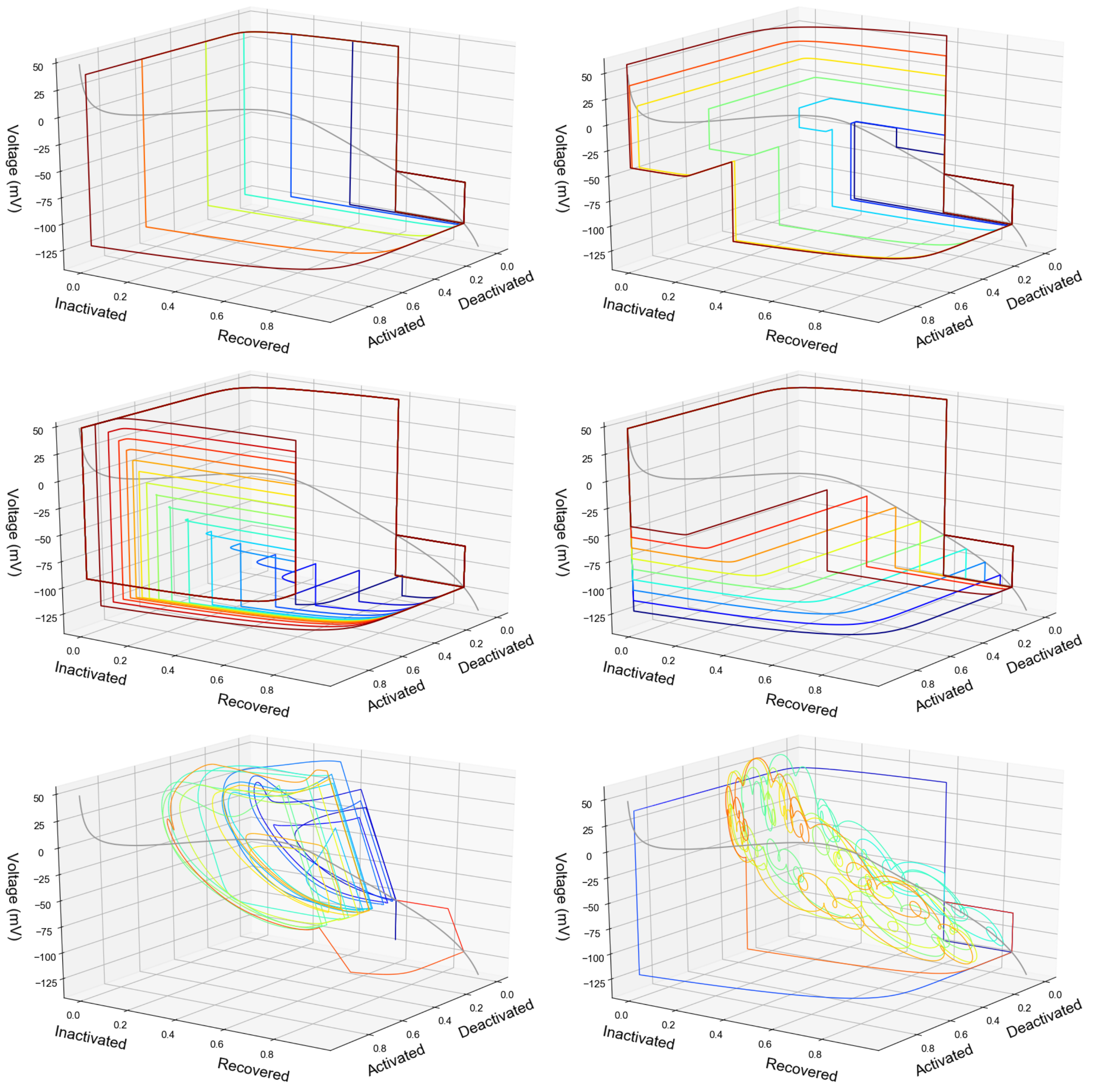

Figure S10: Three-dimensional phase diagrams for Pr2–7.

#### S1.10 Improving experimental protocols

Hints for improved protocol design can be found in the phase plane diagrams shown above. Inspecting the phase plane diagrams in Figure S10, we can see that large parts of Pr2–5 are concerned with setting up the system for a measurement, e.g. waiting to get into a certain steady-state, and subsequently with restoring the original state again. These steps were necessary for manual analysis, but have less use for Method 3. Following the trajectory of the system for these parts of the protocols, we see it is mostly near the x-axis and y-axis. From Figure S1.B we can see that these are areas where the system has low conductance, leading to very small currents and a poor signal-to-noise ratio. For Method 3, these parts provide information about the noise in the signal, but not about the current's dynamics. By contrast, Pr7 spends a large proportion of its time away from axes, in the 'measurable area'.

Looking at the phase planes further, it is tempting to think that exploring the full plane is a desirable property of the protocol. However, our goal in parameterising a model is not to visit every state, but to observe the kinetic parameters ($p_1$ to $p_8$) in action for as many voltages as possible. In other words, we want to observe the current while the system makes each of its four transitions (activation, deactivation, inactivation, and recovery), for all physiologically relevant voltages. (Note that if we had full confidence in our model, a few voltages per parameter would suffice, as the equations constrict the system once a few points are known.) The three-dimensional phase planes shown in Figure S1 demonstrate how the sine wave protocol comes close to realising this ideal. It visits a wide range of physiological voltages, and makes seemingly arbitrary transitions throughout the voltage range. Note however, that it still has relatively low conductance throughout, so that adapting the protocol to start with greater levels of activation may be advantageous. To improve the protocol's performance on predicting deactivation, it may also be useful to add a lower frequency sine wave, causing greater activation while the existing higher frequencies stop the system from inactivating and reducing the amount of observable current.

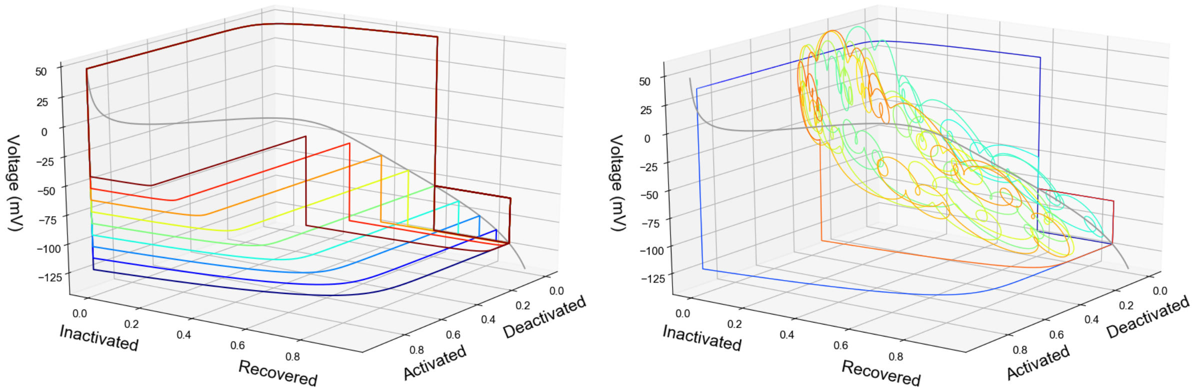

Figure S11: Three-dimensional phase diagrams for Pr5 and Pr7. The relative sparsity of the Pr5 can clearly be seen in this plot, sticking mostly to the walls (where no current is observable) and making only brief controlled forays into observable space. This careful setting up of the right conditions before measuring anything is highly advantageous for traditional analysis, but is not the most efficient strategy for whole-current fitting methods. By contrast the chaotic nature of Pr7 looks very difficult to interpret, but induces all four transitions (activation, deactivation, inactivation, and recovery) at several voltages, all the while producing measurable current.

### S2 SUPPLEMENTARY METHODS

#### S2.1 Experimental data for all 9 cells

Figures showing the data for all 9 cells can be found at https://github.com/CardiacModelling/FourWaysOfFitting.

#### S2.2 Boundaries on the parameter space

Boundaries were defined on the parameter space, based on physiological constraints. This is similar to the concept of a *prior* in Bayesian inference, as it encodes our prior knowledge about the parameter values. As in Beattie et al. (1), we constrained (i) the maximum conductance $p_9 = g_{Kr}$; (ii) the values of the kinetic parameters $p_1$ to $p_8$; and (iii) the reaction rates $k_1$ to $k_4$.

Lower and upper bounds for the maximum conductance in each cell were estimated by Beattie et al. (1), and are shown in Table S1. The lower conductance for each cell was estimated in Beattie et al. (1) by assuming that the current was fully conducting ($a = r = 1$) at some point after the initial +40 mV step of the sine wave protocol (see Beattie et al. (1) for details). An upper bound was then derived by assuming that $a \cdot r > 0.1$ at this point.

Table S1: Cell-specific limits on the conductance parameter

| Cell # | $g_{lower}$ (mS) | $g_{upper}$ (mS) |
|---|---|---|
| 1 | 0.0478 | 0.478 |
| 2 | 0.0255 | 0.255 |
| 3 | 0.0417 | 0.417 |
| 4 | 0.0305 | 0.305 |
| 5 | 0.0612 | 0.612 |
| 6 | 0.0170 | 0.170 |
| 7 | 0.0886 | 0.886 |
| 8 | 0.0434 | 0.434 |
| 9 | 0.0203 | 0.203 |

Bounds for the kinetic parameters were set based on expected physiological ranges of the resulting reaction rates, as well as their expected voltage sensitivity (1):

$$10^{-7}\,\text{ms}^{-1} \leqslant p_i \leqslant 10^{3}\,\text{ms}^{-1}, \quad i \in 1,3,5,7, \tag{S11}$$

$$10^{-7}\,\text{mV}^{-1} \leqslant p_j \leqslant 0.4\,\text{mV}^{-1}, \quad j \in 2,4,6,8. \tag{S12}$$

Additionally, we set lower and upper bounds for the *maximum* transition rates, representing timescales of a minute to a microsecond, using

$$1.67 \cdot 10^{-5}\,\text{ms}^{-1} \leqslant k_i(V = +60) \leqslant 1000\,\text{ms}^{-1}, \quad i \in 1,3, \tag{S13}$$

$$1.67 \cdot 10^{-5}\,\text{ms}^{-1} \leqslant k_j(V = -120) \leqslant 1000\,\text{ms}^{-1}, \quad j \in 2,4. \tag{S14}$$

Here, the values for the lower and upper bounds, again taken from Beattie et al. (1), are chosen to yield (very wide) limits on what can be considered a physiologically realistic maximum reaction rate during the sine wave protocol, which is restricted to a voltage range from −120 mV to +60 mV. The additional rate constraints in Eq. (S13–S14) are functions of two parameters, so they effectively specify 2-dimensional constraints on the parameter pairs $(p_1, p_2)$, $(p_3, p_4)$, $(p_5, p_6)$, and $(p_7, p_8)$. When running optimisations, parameter sets that violated any of the boundary conditions were assigned an error of $\infty$.

### S3 SUPPLEMENTAL RESULTS

#### S3.1 Obtained parameters

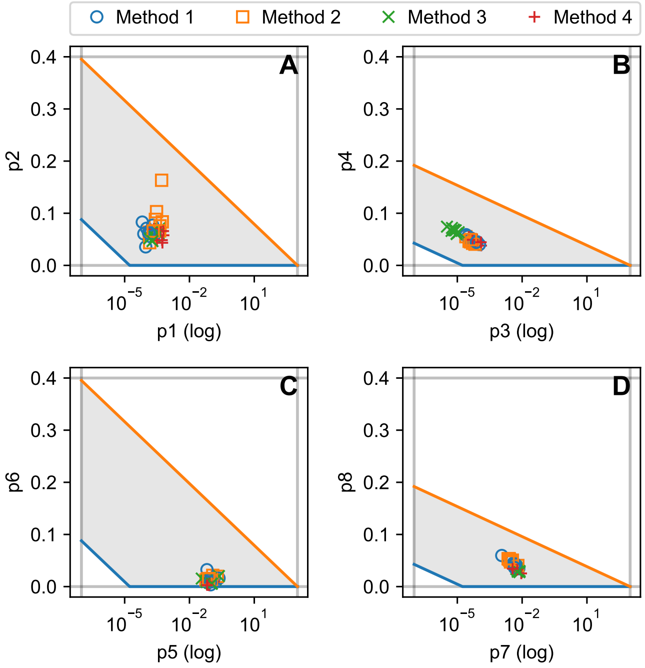

Figure S12: The best parameters returned by all four methods, for all nine cells. Only the eight kinetic parameters are shown. Parameters $p_1$ and $p_2$, shown in panel A, together determine the activation rate, while $p_3$ and $p_4$ in panel B determine deactivation. Similarly, inactivation and recovery are determined by the parameters in panels C and D respectively.

#### S3.2 Validation and cross-validation figures for all cells

Validation and cross-validation figures for all cells can be found at `https://github.com/CardiacModelling/FourWaysOfFitting`.

285 ## S3.3 Relative RMSE tables for all cells

| Cell 1 | Method 1 | Method 2 | Method 3 | Method 4 |
|---|---|---|---|---|
| AP validation | 1.53 | 1.36 | 1.00 | 1.36 |
| Method 1 RMSE | 1.14 | 1.00 | 3.21 | 1.59 |
| Cross-validation M2 | 2.52 | 1.00 | 4.31 | 2.26 |
| Cross-validation M3 | 1.99 | 1.75 | 1.00 | 1.92 |
| Cross-validation M4 | 3.15 | 2.27 | 1.84 | 1.00 |

| Cell 2 | Method 1 | Method 2 | Method 3 | Method 4 |
|---|---|---|---|---|
| AP validation | 2.26 | 2.96 | 1.00 | 1.19 |
| Method 1 RMSE | 1.13 | 1.00 | 2.36 | 1.61 |
| Cross-validation M2 | 2.83 | 1.00 | 3.96 | 2.89 |
| Cross-validation M3 | 2.33 | 2.43 | 1.00 | 2.07 |
| Cross-validation M4 | 3.20 | 4.87 | 1.65 | 1.00 |

| Cell 3 | Method 1 | Method 2 | Method 3 | Method 4 |
|---|---|---|---|---|
| AP validation | 1.14 | 2.10 | 1.00 | 1.37 |
| Method 1 RMSE | 1.00 | 1.07 | 2.85 | 1.94 |
| Cross-validation M2 | 1.83 | 1.00 | 3.48 | 2.18 |
| Cross-validation M3 | 1.29 | 1.46 | 1.00 | 1.54 |
| Cross-validation M4 | 1.66 | 4.92 | 1.77 | 1.00 |

| Cell 4 | Method 1 | Method 2 | Method 3 | Method 4 |
|---|---|---|---|---|
| AP validation | 2.03 | 1.87 | 1.22 | 1.00 |
| Method 1 RMSE | 1.01 | 1.00 | 2.94 | 2.64 |
| Cross-validation M2 | 2.18 | 1.00 | 3.67 | 3.21 |
| Cross-validation M3 | 1.35 | 1.77 | 1.00 | 1.73 |
| Cross-validation M4 | 3.04 | 4.16 | 2.20 | 1.00 |

| Cell 5 | Method 1 | Method 2 | Method 3 | Method 4 |
|---|---|---|---|---|
| AP validation | 1.82 | 4.25 | 1.29 | 1.00 |
| Method 1 RMSE | 1.00 | 1.11 | 4.64 | 1.99 |
| Cross-validation M2 | 2.58 | 1.00 | 6.33 | 2.48 |
| Cross-validation M3 | 2.46 | 3.88 | 1.00 | 2.13 |
| Cross-validation M4 | 3.78 | 13.54 | 1.70 | 1.00 |

| Cell 6 | Method 1 | Method 2 | Method 3 | Method 4 |
|---|---|---|---|---|
| AP validation | 2.92 | 1.00 | 1.07 | 1.31 |
| Method 1 RMSE | 2.32 | 1.00 | 3.74 | 2.05 |
| Cross-validation M2 | 4.48 | 1.00 | 5.20 | 3.17 |
| Cross-validation M3 | 2.41 | 1.79 | 1.00 | 2.29 |
| Cross-validation M4 | 3.82 | 2.13 | 1.60 | 1.00 |

| Cell 7 | Method 1 | Method 2 | Method 3 | Method 4 |
|---|---|---|---|---|
| AP validation | 2.04 | 5.54 | 1.12 | 1.00 |
| Method 1 RMSE | 1.00 | 1.27 | 5.85 | 3.03 |
| Cross-validation M2 | 1.56 | 1.00 | 4.71 | 2.36 |
| Cross-validation M3 | 1.36 | 2.51 | 1.00 | 1.47 |
| Cross-validation M4 | 2.71 | 6.88 | 2.09 | 1.00 |

| Cell 8 | Method 1 | Method 2 | Method 3 | Method 4 |
|---|---|---|---|---|
| AP validation | 2.09 | 1.56 | 1.00 | 1.09 |
| Method 1 RMSE | 1.35 | 1.00 | 4.79 | 2.28 |
| Cross-validation M2 | 2.64 | 1.00 | 7.07 | 2.87 |
| Cross-validation M3 | 1.61 | 1.50 | 1.00 | 1.47 |
| Cross-validation M4 | 2.90 | 2.35 | 1.93 | 1.00 |

| Cell 9 | Method 1 | Method 2 | Method 3 | Method 4 |
|---|---|---|---|---|
| AP validation | 1.42 | 1.71 | 1.00 | 1.25 |
| Method 1 RMSE | 1.00 | 1.01 | 2.21 | 1.81 |
| Cross-validation M2 | 2.34 | 1.00 | 3.36 | 3.59 |
| Cross-validation M3 | 1.57 | 1.61 | 1.00 | 1.52 |
| Cross-validation M4 | 1.66 | 1.94 | 1.43 | 1.00 |

| All cells | Method 1 | Method 2 | Method 3 | Method 4 |
|---|---|---|---|---|
| AP validation | 1.7 (0.3) | 2.2 (1.1) | 1.0 (0.1) | 1.1 (0.2) |
| Method 1 RMSE | 1.2 (0.5) | 1.0 (0.2) | 3.3 (0.6) | 1.9 (0.2) |
| Cross-validation M2 | 2.5 (0.7) | 1.0 (0.1) | 4.6 (1.0) | 2.8 (0.4) |
| Cross-validation M3 | 1.7 (0.4) | 2.0 (0.9) | 1.0 (0.4) | 1.7 (0.4) |
| Cross-validation M4 | 2.8 (0.9) | 4.3 (2.2) | 1.8 (0.5) | 1.0 (0.3) |

Figure S13: Relative RMSE for all cells.

#### S3.4 Performance

Figure S14 shows the experimental and computational effort that goes each method. Performing experiments for analysis with Method 4 is considerably faster than the experiments needed for methods 1–3, with only 8 seconds needed compared to 228 s for the Pr2–5 combination. This long protocol duration also has an effect on the time needed to run numerical optimisations. The time for a single optimisation (one of the fifty repeats we ran per method) is shown for methods 2–4 in the top right panel. Method 1 is deterministic and runs almost instantaneously, so is omitted here. Method 2 takes far longer than the other methods, and Method 3 is slower than Method 4. Inspecting the number of evaluations (simulations) performed by each method, we see that Method 2 is much more similar to methods 3 and 4 in this respect. A large number of Method 2 optimisations terminated with a low number of evaluations, which may indicate they stopped exploring early and hit a local minimum.

Since the difference in time per optimisation between methods 3 and 4 is not explained by the number of evaluations, it must be due to the time per evaluation (simulation). The final panel shows that this is indeed much larger for Method 4 than Method 3. The time per evaluation for Method 2 is larger still, since Method 2 not only runs a simulation every evaluation, but also needs to post-process the results to extract time constants and steady states.

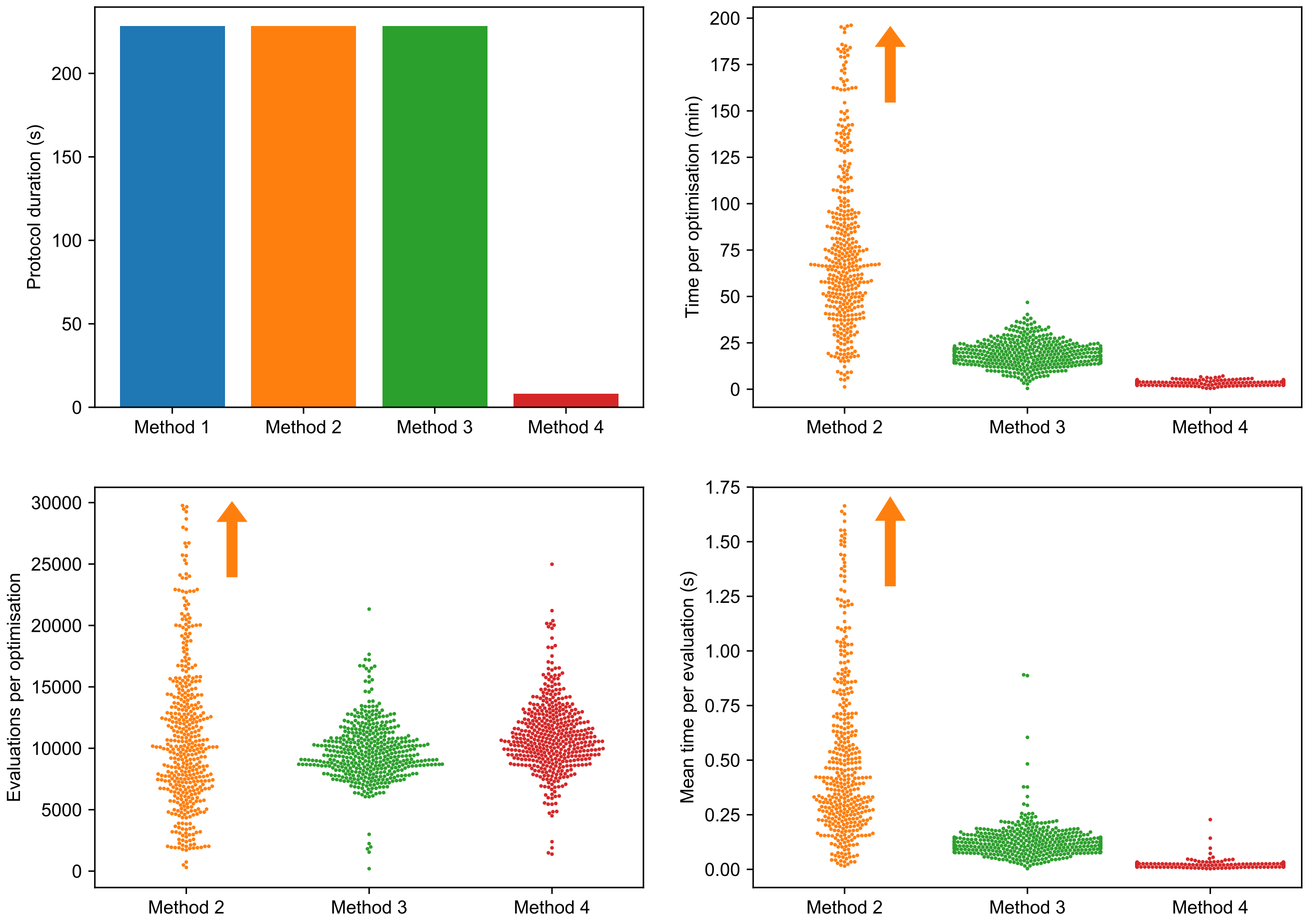

Figure S14: Experimental duration and optimisation duration. (*Top-left*) The duration of the protocols needed for each method: 228s for methods 1–3 (which are based on Pr2–5), and 8 s for Method 4 (based on Pr7). (*Top-right*) The time taken for a single optimisation, with 50 points shown per method per cell. (*Bottom-left*) The number of evaluations per optimisation for methods 2–4. (*Bottom-right*) The mean time per evaluation, calculated for 50 optimisations for each cell. Orange arrows indicate where the $y$-axis is hiding some outliers from Method 2.

#### S3.5 Cross-sections of the optimisation surfaces

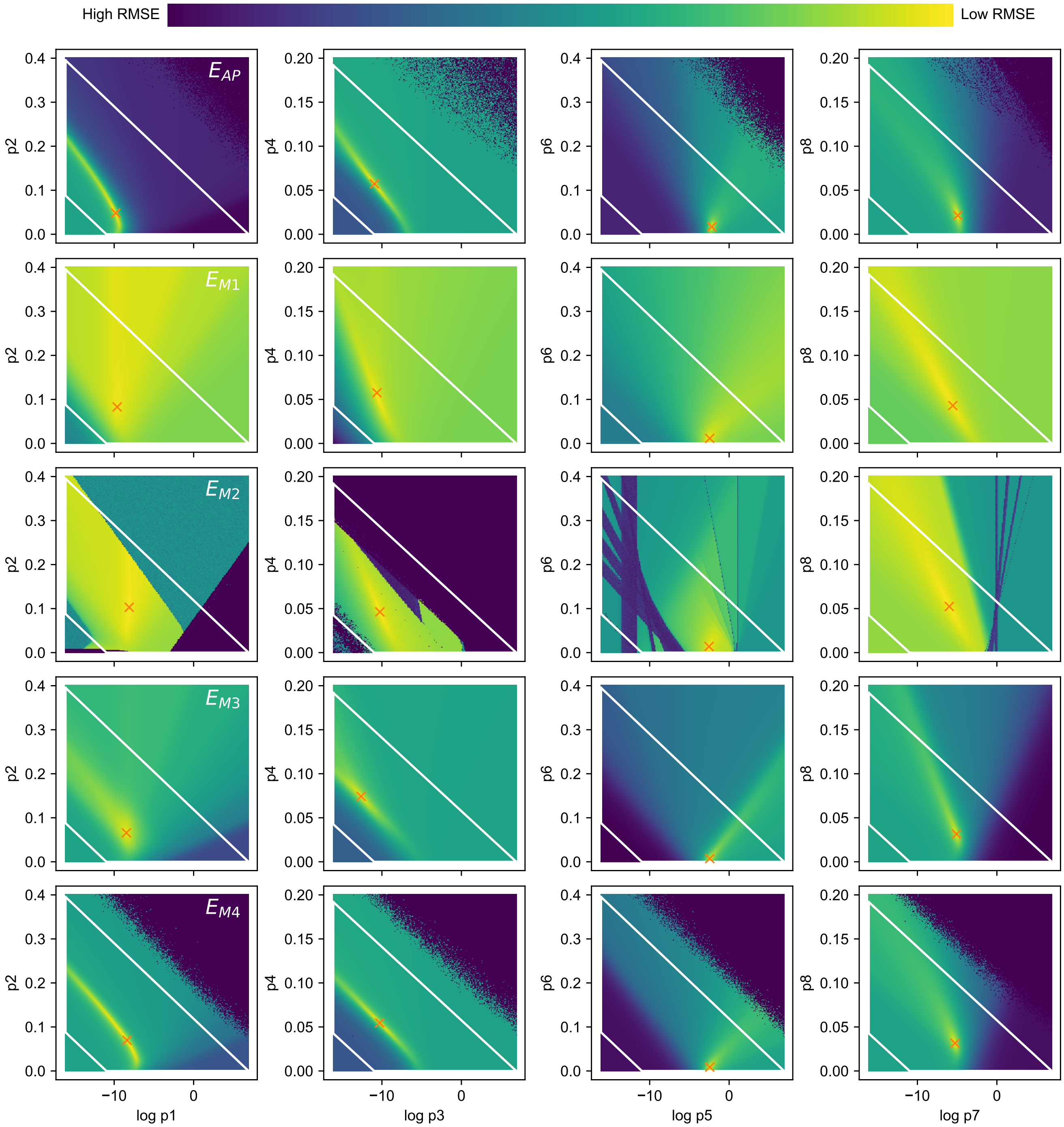

Figure S15: Cross-sections of all optimisation criteria for Cell #5, for $E_{\mathrm{AP}}$ (*Top*), and $E_1$–$E_4$. The surfaces were drawn by performing a brute-force mapping (256x256 evaluations) around a fixed point. For $E_{\mathrm{AP}}$ this fixed point was chosen by first running an optimisation to find its optimum. For $E_1$–$E_4$ the fixed point was the result returned by methods 1–4. Note that for $E_1$ this point does not correspond exactly to $E_1$'s minimum.

#### S3.6 Method 1b: Minimising $E_{\mathrm{M1}}$

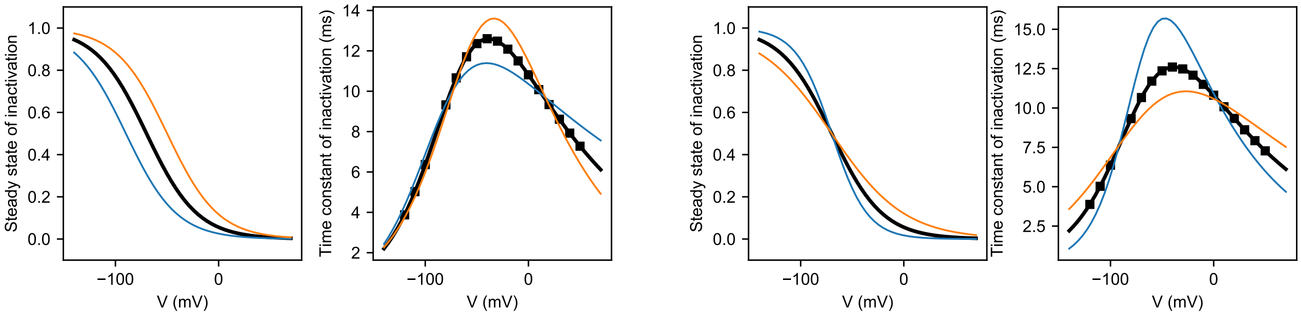

Figure S16: In Method 1, the steady-state approximations are used in deriving the approximations of the time constants. The time constants are relatively robust against shifts in the midpoints of (in)activation, but react more strongly to changes in the steady state slopes. For Method 1, this implies that a small error in estimating the slope (for example do to having points near the reversal potential) can cause a large error in the time constants. For Method 2, it shows that a better fit on the time constants can be obtained by slightly tweaking the steady state curves.

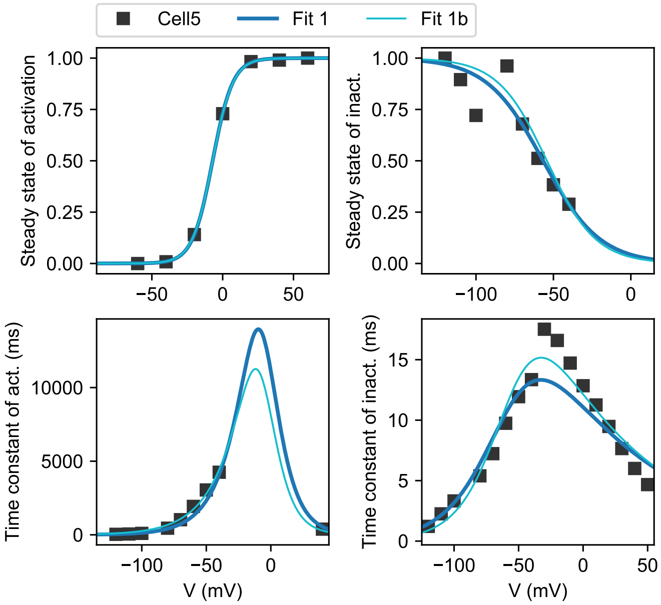

Figure S17: Minimising $E_{\mathrm{M1}}$ directly allows for a trade-off between goodness-of-fit in the steady states and the time constants. This results in improved time-constant fits, without much change to the steady-state fits. Data is shown for Cell #5.

| Cell 5 | Method 1 | Method 2 | Method 3 | Method 4 |
|---|---|---|---|---|
| AP validation | 1.82 | 4.25 | 1.29 | 1.00 |
| Method 1 RMSE | 1.00 | 1.11 | 4.64 | 1.99 |
| Cross-validation M2 | 2.58 | 1.00 | 6.33 | 2.48 |
| Cross-validation M3 | 2.46 | 3.88 | 1.00 | 2.13 |
| Cross-validation M4 | 3.78 | 13.54 | 1.70 | 1.00 |

| Cell 5 | Method 1b | Method 2 | Method 3 | Method 4 |
|---|---|---|---|---|
| AP validation | 1.68 | 4.25 | 1.29 | 1.00 |
| Method 1 RMSE | 1.00 | 1.43 | 5.96 | 2.56 |
| Cross-validation M2 | 1.90 | 1.00 | 6.33 | 2.48 |
| Cross-validation M3 | 3.32 | 3.88 | 1.00 | 2.13 |
| Cross-validation M4 | 5.89 | 13.54 | 1.70 | 1.00 |

Figure S18: Using Method 1b (*Right*) leads to E1 RMSEs that outperform Method 1 (*Left*). However, this improvement does not necessarily translate to better predictions, as the fundamental idea — that the model equations should be overlaid on their experimental approximates — is still flawed. Data is shown for Cell #5.

##### S3.7 Method 2b: Minimising $E_{\mathrm{M2}}$ starting from Method 1 result

A Method 2 variant (“Method 2b”) can be created by adapting Method 2 to only perform a single run, starting from the parameters returned by Method 1. In the experiments we ran, this gave very similar results to Method 2, but at a lower computational costs. However, these results will only generalise if the function $E_{\mathrm{M2}}$ is smooth and easy to navigate between the Method 1 and Method 2 results.

Table S2: Method 2 and Method 2b results

| Cell | $E_{\mathrm{M2}}$ Method 2 | $E_{\mathrm{M2}}$ Method 2b |
|---|---|---|
| Cell | Method 2 | Method 2b |
| 1 | 0.1852969815 | 0.1900501636 |
| 2 | 0.1807762308 | 0.1822721183 |
| 3 | 0.2155275628 | 0.2155275628 |
| 4 | 0.1648576405 | 0.1685394193 |
| 5 | 0.1583982474 | 0.1583982474 |
| 6 | 0.1698205828 | 0.1669494977 |
| 7 | 0.1335628150 | 0.1338667076 |
| 8 | 0.1654924563 | 0.1654924563 |
| 9 | 0.1848795472 | 0.1900065180 |

##### S3.8 Method 3b: Minimising $E_{\mathrm{M2}}$ starting from Method 1 result

A similar adaptation can be used to create a Method 3 variant, “Method 3b”.

Table S3: Method 3 and Method 3b results

| Cell | $E_{\mathrm{M3}}$ Method 3 | $E_{\mathrm{M3}}$ Method 3b |
|---|---|---|
| 1 | 0.0448121214 | 0.0448121214 |
| 2 | 0.0479263275 | 0.0479263275 |
| 3 | 0.0613629021 | 0.0613629021 |
| 4 | 0.0801896081 | 0.0801896081 |
| 5 | 0.0394435210 | 0.0394435210 |
| 6 | 0.0666019884 | 0.0666019884 |
| 7 | 0.1164911334 | 0.1164911334 |
| 8 | 0.1005265480 | 0.1005265480 |
| 9 | 0.0683642823 | 0.0683642823 |
